# Antibiotic resistome and microbial community structure during anaerobic co-digestion of food waste, paper and cardboard

**DOI:** 10.1101/564823

**Authors:** Kärt Kanger, Nigel G.H. Guilford, HyunWoo Lee, Camilla L. Nesbø, Jaak Truu, Elizabeth A. Edwards

## Abstract

Antimicrobial resistance is a globally recognized public health risk. High incidence of antibiotic resistant bacteria and antibiotic resistance genes (ARGs) in solid organic waste necessitates the development of effective treatment strategies. The objective of this study was to assess ARG diversity and abundance as well as the relationship between resistome and microbial community structure during anaerobic co-digestion (AD) of food waste, paper and cardboard. A lab-scale solid-state AD system consisting of six sequentially fed leach beds (each with a solids retention time of 42 days) and an upflow anaerobic sludge blanket (UASB) reactor was operated under mesophilic conditions continuously for 88 weeks to successfully treat municipal organic waste and produce biogas. A total of ten samples from digester feed and digestion products were collected for microbial community analysis including SSU rRNA gene sequencing, total community metagenome sequencing and quantitative PCR. Taxonomic analyses revealed that AD changed the taxonomic profile of the microbial community: digester feed was dominated by bacterial and eukaryotic taxa while anaerobic digestate possessed a large proportion of archaea mainly belonging to the methanogenic genus *Methanosaeta*. ARGs were identified in all samples with significantly higher richness and relative abundance per 16S rRNA gene in digester feed compared to digestion products. Multidrug resistance was the most abundant ARG type. AD was not able to completely remove ARGs as shown by ARGs detected in digestion products. Using metagenomic assembly and binning we detected potential bacterial hosts of ARGs in digester feed, that included *Erwinia, Bifidobacteriaceae, Lactococcus lactis* and *Lactobacillus*.

**IMPORTANCE:** Solid organic waste is a significant source of antibiotic resistance genes (ARGs) (1) and effective treatment strategies are urgently required to limit the spread of antimicrobial resistance. Here we studied the antibiotic resistome and microbial community structure within an anaerobic digester treating a mixture of food waste, paper and cardboard. We observed a significant shift in microbial community composition and a reduction in ARG diversity and abundance after 6 weeks of digestion. We identified the host organisms of some of the ARGs including potentially pathogenic as well as non-pathogenic bacteria, and we detected mobile genetic elements required for horizontal gene transfer. Our results indicate that the process of sequential solid-state anaerobic digestion of food waste, paper and cardboard tested herein provides a significant reduction in the relative abundance of ARGs per 16S rRNA gene.

## INTRODUCTION

Antimicrobial resistance (AMR) is a widely recognized public health risk. The extensive use of antimicrobial compounds since World War II has triggered the rapid spread and evolution of AMR mechanisms to such an extent that AMR is considered one of today’s top medical concerns globally (2). Currently, 700 000 deaths are attributed to resistant microbial infections annually, which is projected to increase to 10 million deaths by 2050 resulting in the loss of 100 trillion USD of economic output (3). In light of this gloomy prospect, a One-Health approach has been proposed to tackle AMR by recognizing the connections between human and animal health and the environment (4).

Although the vast majority of antimicrobial research is focusing on the clinical setting, the natural environment has gained attention as a possible reservoir and dispersal route of antimicrobial resistance determinants, including antibiotic resistant bacteria and antibiotic resistance genes (ARGs) (5, 6). Various natural ecosystems, such as ancient permafrost sediments (7), pristine soil (8, 9), oceanic (10) and freshwater bodies (11) have been shown to possess a diversity of ARGs collectively defined as the resistome. However, higher concentrations of resistance determinants have been observed in environments with anthropogenic impact where selective pressure for ARGs is present, such as wastewater treatment plants, animal husbandry facilities, aquaculture farms and pharmaceutical manufacturing (12).

Various types of solid organic waste may also serve as potential sources of antibiotic resistant bacteria and ARGs (1). In addition to the extensively studied resistomes of sewage sludge and animal manure, the organic fraction of municipal solid waste (MSW) may also contribute to the dissemination of AMR. Global production of MSW is estimated at around 2 billion tons per year, of which 34–53% is organic biodegradable waste including food waste as the primary component (13). Paper waste forms 17% of MSW globally, including different lignocellulosic fibres such as cardboard, newspaper, magazines, wrapping paper, shredded paper, boxes, bags and beverage cups (14).

Several studies have highlighted the role of the food chain in AMR as a direct link to human health (15, 16) and identified antibiotic resistant pathogens in various food products such as meat (17–19), fruit and vegetables (20–23), poultry (24, 25), fish (26, 27) and dairy (28). Therefore, it is reasonable to assume that ARGs are also present in food waste: Lee *et al*. (29) detected a variety of ARGs in the wastewater from food waste recycling in Korea, although the abundance of ARGs remained below that of manure and sewage sludge.

To mitigate the harmful environmental impacts of organic solid waste, anaerobic digestion (AD) is widely implemented as a treatment strategy, providing waste stabilization as well as the production of renewable energy (30). AD changes the structure of the microbial community of the substrates which in turn affects the resistome present in digestion products. Although it is generally accepted that ARG abundance is reduced overall during AD, enrichment of some ARGs and rebound-effects have been reported in several studies (summarized by Youngquist *et al*. (31)). For example, Pu *et al*. (32) studied the impact of applying pig manure to fields and found that AD reduced the relative abundance of Macrolide-Lincosamide-Streptogramin (MLS) and tetracycline resistance genes, while resistance genes for sulfa, aminoglycoside, florfenicol, amphenicol and chloramphenicol were enriched. Similarly, the effect of aerobic composting of organic waste remains contradictory: while some evidence suggests composting can reduce the abundance of antibiotic resistant bacteria and ARGs, other studies have shown increased abundance and diversity of ARGs (31).

Recently, there has been growing interest in AD of food waste due to its high energetic value (13). However, limited information is available on the effect of AD on the microbial community and resistome in food waste. Zhang *et al*. (33) identified 11 ARGs and class 1 integron integrase gene *intI1* in digested food waste: while *tetA, tetB, tetX, sul1, cmlA, floR* and *intI1* were significantly reduced by AD, enrichment of *tetM, tetW, tetQ* and *tetO* was recorded. Similarly, another study detected both increase and reduction of specific ARGs in the co-digestion of sewage sludge and food waste following microwave pretreatment (34). Thus, the effect of AD on the food waste resistome and the associated microbial community is still unclear.

In addition to quantification of individual ARGs, detecting potential host organisms of these genes is important to evaluate the risk to human health. Several studies have used correlation and network analysis to detect relationships between individual ARGs and bacterial genera (34–37). High mobility of ARGs due to horizontal gene transfer (HGT) is responsible for the spread of AMR between different bacteria including human pathogens as well as non-pathogenic environmental bacteria which often serve as reservoirs of ARGs (38).

The objective of this study was to measure ARG abundance and resistome diversity before and after anaerobic co-digestion of food waste, paper and cardboard. Samples were collected from a lab-scale solid-state AD system that exhibited stable methane production and substrate destruction rates (39, 40). By simultaneously analyzing microbial community shifts, we attempted to assign ARGs to specific host organisms. Using SSU rRNA gene sequencing, total community metagenome sequencing and targeted quantitative PCR we were able to quantify changes in the diversity and size of the resistome before and after digestion together with changes in the taxonomic profile of the microbial community. We were further able to identify potential host organisms of ARGs by using metagenomic assembly and binning methods.

## RESULTS

A total of 10 samples were collected from a lab-scale solid-state anaerobic digestion system described in detail by a recent PhD thesis (39). The ten samples included three samples of raw food waste (FW1, FW2, FW3) collected from local residential green bin program, three samples of the mixture of food waste and lignocellulosic fibres that constituted the leach bed feed (LBF1, LBF2, LBF3), three samples of 6 week old digestate (DG74, DG76, DG78) and one sample of the microbial granules from the upflow anaerobic sludge blanket reactor (UASB) treating leachate from the leach beds. DNA was extracted from each sample in order to analyze the microbial community composition and ARGs diversity, abundance, genomic location and host organisms using a multi-pronged approach combining metagenome sequence analyses, taxonomic profiling and quantitative PCR (Fig. 1).

**FIG 1.**
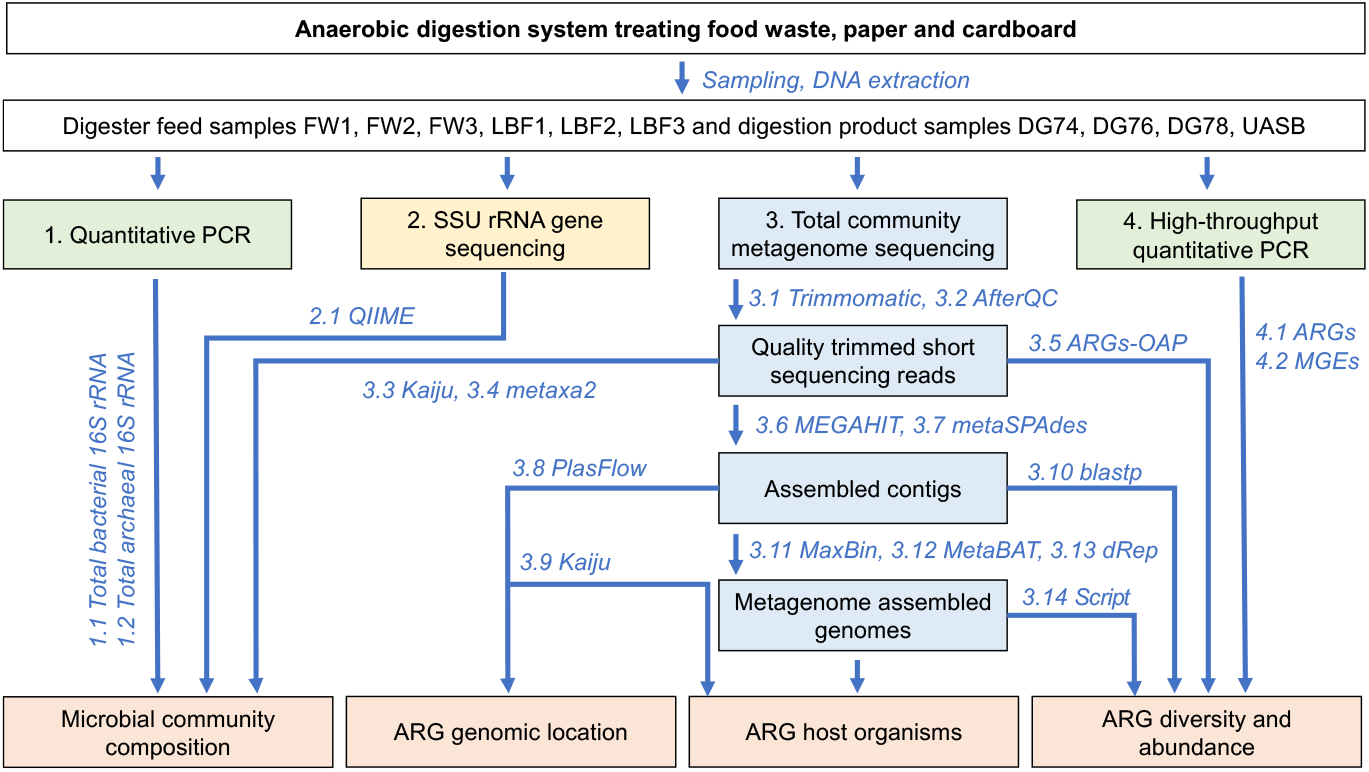
Experimental workflow of anaerobic digestion experiment. Total community DNA extracts from 10 samples including food waste (FW1, FW2, FW3), leach bed feed (LBF1, LBF2, LBF3), digestate (DG74, DG76, DG78) and microbial granules from UASB reactor (UASB) were subjected to small subunit (SSU) rRNA gene amplicon sequencing (2), total community metagenome sequencing (3) and quantitative PCR analysis (1, 4). Sequencing data was analyzed using a variety of bioinformatic tools shown and numbered in italics. SSU rRNA gene sequencing data was analyzed using QIIME1 (tool 2.1). The workflow for total community metagenome sequencing data included quality trimming of the short sequencing reads (tools 3.1 and 3.2), taxonomic annotation of the quality trimmed reads (tools 3.3 and 3.4), detection of ARGs from quality trimmed reads (tool 3.5), assembly of the quality trimmed reads into contigs (tools 3.6 and 3.7), identification of plasmid sequences (tool 3.8) and ARGs (tool 3.10) from assembled contigs, taxonomic annotation of assembled contigs (tool 3.9), binning of the assembled contigs into metagenome assembled genomes (tools 3.11, 3.12, 3.13) and detection of ARGs in metagenome assembled genomes by a local Python script (tool 3.14). Quantitative PCR was performed for bacterial (1.1) and archaeal (1.2) 16S rRNA gene as well as for ARGs (4.1) and MGEs (4.2) using a high-throughput qPCR array. Results covered microbial community composition, ARG diversity, abundance, genomic location and potential bacterial host organisms of ARGs.

### Microbial community

Microbial community structure was analyzed by small subunit (SSU) rRNA gene-fragment sequencing (method 2 in Fig. 1) capturing bacterial, archaeal and eukaryotic diversity (Tables S1 and S2 in supplemental material). A clear distinction in the communities of digester feed and digestion products was detected: food waste and leach bed feed microbial communities were dominated by bacterial and eukaryotic taxa while anaerobic digestate and microbial community of granules from the UASB reactor possessed a large proportion of archaeal phylotypes mainly belonging to the methanogenic genus *Methanosaeta* (Fig. 2). The most abundant bacterial OTUs in digester feed belonged to the family *Enterobacteriaceae*, including the genera *Citrobacter, Enterobacter, Erwinia, Kluyvera, Pantoea, Serratia* and *Lelliottia*. Additionally, representatives of the phylum *Firmicutes* were detected in FW2, including the genera *Lactobacillus* and *Leuconostoc*, probably indicating fermentation processes occurring in the respective food waste. In addition to bacterial sequences, fungal and plant material was detected in digester feed (FW, LBF), but not in digestion products (DG) or the UASB. Across all samples, 36–62% of OTUs were detected at low relative abundances comprising less than 1% of the total community. A comparison of genus level taxonomic composition detected from SSU rRNA gene amplicon sequence analysis using QIIME (tool 2.1 in Fig. 1) to that derived from total community metagenome sequencing analysis using Kaiju (tool 3.3 in Fig. 1) or metaxa2 (tool 3.4 in Fig. 1) showed relatively good agreement between the three different annotation tools used in this study (Fig. S1 in supplemental material).

**FIG 2.**
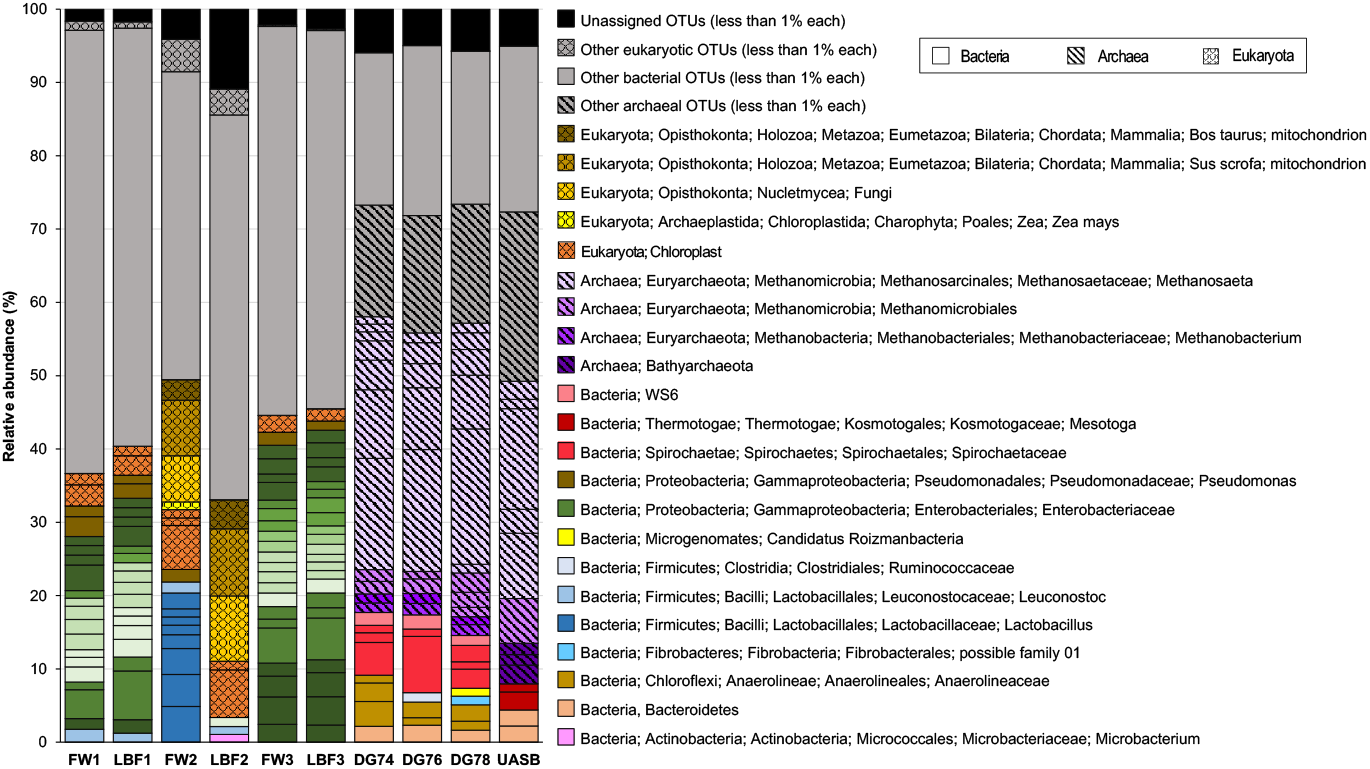
Taxonomic composition of bacterial, archaeal and eukaryotic OTUs detected by SSU rRNA gene sequencing. Each rectangle represents an individual OTU with relative abundance ≥1%. The remaining OTUs with relative abundance <1% are summed and represented in grey as separate rectangles for bacteria, archaea and eukaryota. Bacterial OTUs are presented in solid colors, archaeal OTUs with vertical lines and eukaryotic OTUs in diamond pattern. OTUs belonging to the bacterial family *Enterobacteriaceae* are presented in various shades of green. Additional information on all detected OTUs is provided in Table S2 in supplemental material.

### ARG diversity and abundance

Total community metagenome sequencing data from seven samples (Table S3 in supplemental material) was analyzed using the ARGs Online Analysis Pipeline (ARGs-OAP) (41) (tool 3.5 in Fig. 1) to characterize the distribution and diversity of ARGs before and after anaerobic digestion. The richness of ARGs, measured as the number of distinct ARGs identified in one sample type, was highest in digester feed with 330 and 336 different ARGs detected in FW and LBF samples, respectively. The richness of ARGs in digestion products remained two times lower with 115 different ARGs detected in DG and UASB samples indicating reduced diversity of ARGs after anaerobic digestion. Twenty three ARGs were found to be unique to digestion products belonging to aminoglycoside (*aac(3)-I*, *ant(9)-I*), beta-lactam (*OXA-10, OXA-205, OXA-251, OXA-34, OXA-46, OXA-75*), chloramphenicol (*catQ*), MLS (*ereB, lnuB, mphA, carA*), sulfonamide (*sul3*), tetracycline (*tet44, tetT*), trimethoprim (*dfrA5*) and vancomycin (*vanA, vanG, vanH, vanN, vanU, vanX*) type.

In addition to richness, the relative abundance of ARGs per 16S rRNA gene was determined from total community metagenome sequencing data (Table S4 in supplemental material). Relative abundances of ARGs per 16S rRNA gene were significantly higher in digester feed than in digestion products, indicating the ability of anaerobic digestion to reduce ARGs (Fig. 3). Total relative abundances of ARGs per 16S rRNA gene in food waste samples FW1 (0.78 ARG/16S rRNA) and FW2 (0.40 ARG/16S rRNA) were similar to the respective values in the mixtures of food waste with lignocellulosic fibres (LBF1 0.84 ARG/16S rRNA, LBF2 0.44 ARG/16S rRNA), indicating the role of food waste as the primary source of ARGs in leach bed feed. Most of the ARGs identified in digester feed conferred multidrug resistance (Fig. 3A). Additionally, many potential ARGs with unclassified resistance type were detected. The highest relative abundance value for an individual gene (Fig. 3B) was recorded for AcrAB-TolC multidrug efflux complex subunit *acrB* reaching 0.05 copies/16S rRNA gene in samples FW1 and LBF1. Other highly abundant genes in digester feed included subunits of multidrug resistance complexes (acrA, *mdtB, mdtC, tolC*), several regulatory protein genes associated with AMR (*cpxR, arlR*) and bacitracin resistance gene *bacA*. Relative abundances of individual ARGs in digestion products remained below 0.01 copies/16S rRNA in all cases. Similarly to digester feed, the highest value for an individual gene in digestion products was recorded for *acrB* with 0.007 copies/16S rRNA in UASB.

**FIG 3.**
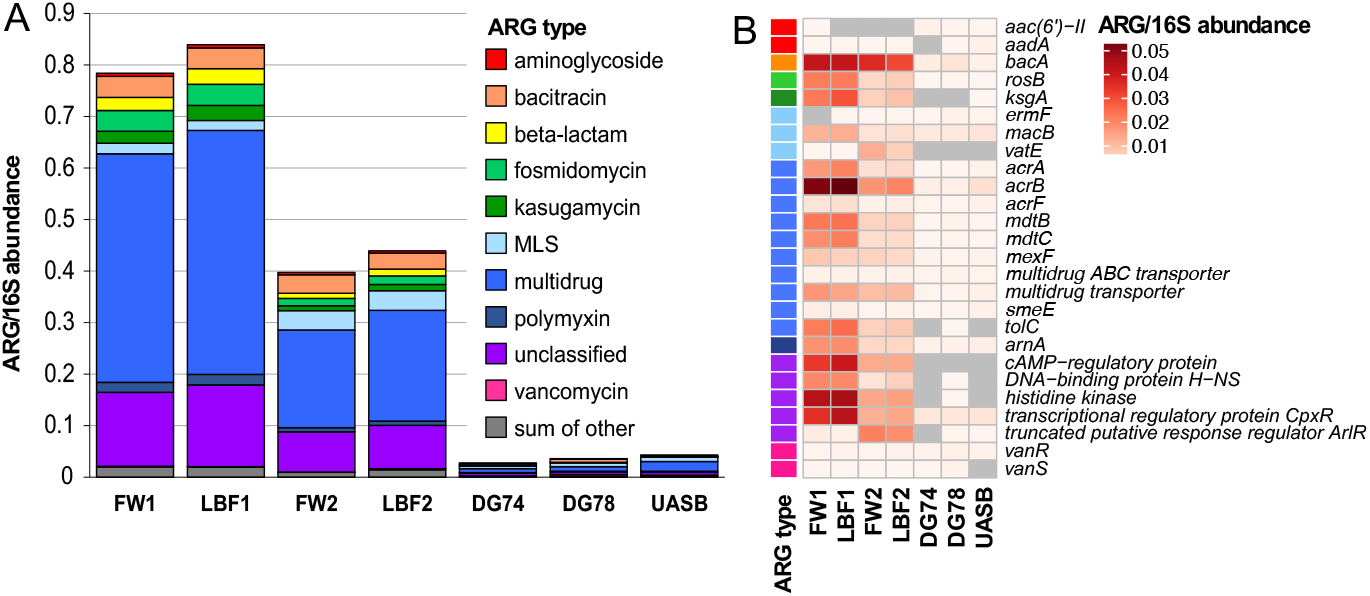
Relative abundance of ARGs per 16S rRNA gene categorized by type (A) and by gene (B) detected in total community metagenome sequencing reads using ARGs-OAP. The ten most abundant ARGs from each sample are depicted in panel B. Genes not detected are marked in grey. Relative abundances of all detected ARGs are provided in Table S4 in supplemental material.

A high-throughput qPCR (HT-qPCR) array was also used as an alternative approach to quantify 315 ARGs (tool 4.1 in Fig. 1) and 57 mobile genetic elements (MGEs) (tool 4.2 in Fig. 1) in food waste and digestate DNA samples (Table S5 in supplemental material). In agreement with the results from the metagenomic data, the diversity of ARGs and MGEs was higher in food waste with 161 different genes detected, while only 32 different genes were detected in digestate (Fig. 4). Among these, 10 ARGs were found only in digestate samples and not in food waste. Notably, MLS resistance genes *mphA* and *lnuB*, phenicol resistance gene *catQ* and tetracycline resistance gene *tet44* were found to be unique to digestate by both methods.

**FIG 4.**
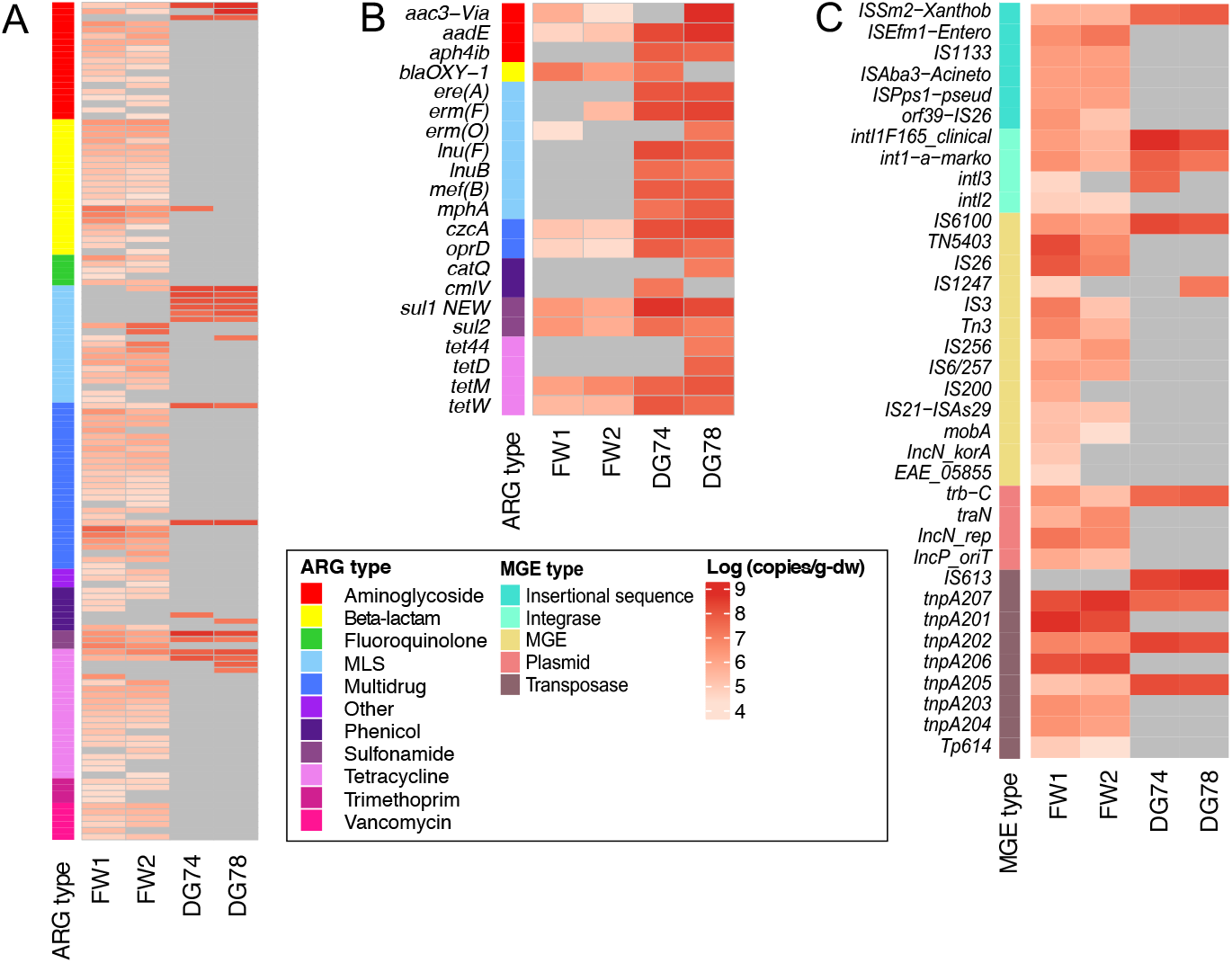
Absolute abundance of ARGs and MGEs in food waste (FW1, FW2) and digestate (DG74, DG78) samples detected using a high-throughput qPCR array. Absolute abundance of all detected genes is presented on log-scale as gene copies per gram of dry weight of food waste or digestate samples. Genes not detected are marked in grey. Panel A shows all ARGs detected in at least one of the four samples. Panel B shows all ARGs detected in at least one of the digestate samples. Panel C shows all MGEs detected in at least one of the four samples. Raw data for all analyzed genes together with abundance calculations are provided in Table S5 in supplemental material.

Although the richness of ARGs and MGEs was higher in food waste samples, absolute abundances of individual genes normalized per sample dry weight reached higher levels in digestates (Fig. 4) according to HT-qPCR array results calculated based on abundance of 16S rRNA gene per gram of dry sample (tool 1.1 in Fig. 1, Fig. S2 in supplemental material). Most prevalent ARG types in final digestates (Fig. 4B) included MLS (genes *ere(A), erm(F), erm(O), lnu(F), lnuB, mef(B), mphA*), aminoglycoside (genes *aac3-Via, aadE, aph4ib*) and tetracycline (genes *tet44, tetD, tetM, tetW*) with absolute abundances of individual genes ranging between 10^7^ and 10^8^ copies/g-dw. Absolute abundances of ARGs in food waste samples ranged between 10^4^ and 10^7^ copies/g-dw. MGEs followed a similar pattern to ARGs with more genes detected in food waste but higher absolute abundances of individual genes per gram of sample dry weight detected in digestates (Fig. 4C).

### Genomic context of ARGs

In order to study the genomic context of ARGs, total community metagenome sequencing reads were assembled into longer contigs using MEGAHIT (tool 3.6 in Fig. 1) and metaSPAdes (tool 3.7 in Fig. 1) assemblers (Table S6 in supplemental material), followed by detection of ARGs on contigs (tool 3.10 in Fig. 1) and identification of plasmid sequences carrying ARGs (tool 3.8 in Fig. 1) (Table S7 in supplemental material).

On average, 0.10% of all assembled contigs carried ARGs in digester feed, while only 0.003% of contigs identified in digestion products included ARGs (Table S8 in supplemental material). More than 90% of ARG-carrying contigs in all samples included only one ARG, however, contigs with multiple ARGs were also observed with up to five ARGs per contig (Table S8 in supplemental material) detected in digester feed. Contigs with multiple ARGs typically carried subunits of multidrug efflux systems such as MdtABC-TolC coupled with two-component regulatory systems for efflux proteins (such as BaeSR).

Identification of plasmids from ARG-carrying contigs revealed that on average 32% of ARGs detected in digestion substrates were located on plasmids (Table 1) indicating their potential for HGT. The distribution of plasmid ARGs resembled the pattern of ARGs identified from short sequencing reads with multidrug resistance genes being the most abundant type in digestion substrates (Fig. S3 in supplemental material). Proportion of plasmid ARGs in digestion products varied between 33.3% and 60.0%, although the number of detected ARGs remained low (11 ARGs on plasmids in DG74, 24 ARGs in DG78, and 20 ARGs in UASB, respectively; Table 1). ARGs identified as chromosomal accounted for 8.3 to 28.9% of ARGs across all samples, remaining below the respective values for plasmid ARGs in all cases (Table 1).

**TABLE 1.** Genomic location of antibiotic resistance genes (ARGs) detected in total community metagenome sequencing data assembled with MEGAHIT. ARGs were detected by BLAST analysis of protein coding genes using local SARG reference database. Plasmid or chromosomal location of the ARGs was predicted by PlasFlow. All ARG BLAST hits detected in MEGAHIT-assembled metagenome sequencing data are provided in Table S7 in supplemental material.

| Sample | Total assembly length (bp) | Protein coding genes | Number of ARG BLAST hits (% from protein coding genes) | Plasmid ARGs (%) | Chromosomal ARGs (%) |
| --- | --- | --- | --- | --- | --- |
| FW1 | 1 285 684 727 | 2 088 107 | 2 407 (0.12%) | 30.2 | 28.6 |
| FW2 | 941 870 975 | 1 796 492 | 2 484 (0.14%) | 34.7 | 23.0 |
| LBF1 | 120 537 563 | 1 821 640 | 1 908 (0.10%) | 30.0 | 28.9 |
| LBF2 | 1 044 915 087 | 1 906 512 | 1 938 (0.10%) | 33.2 | 21.9 |
| DG74 | 726 434 840 | 1 137 099 | 11 (0.001%) | 36.4 | 9.1 |
| DG78 | 729 045 788 | 1 138 132 | 24 (0.002%) | 33.3 | 8.3 |
| UASB | 941 623 397 | 1 636 065 | 20 (0.001%) | 60.0 | 10.0 |

### Bacterial hosts of ARGs

MEGAHIT-assembled contigs that were not identified as plasmids were further annotated for their taxonomic affiliation (tool 3.9 in Fig. 1). Figure 5 depicts the frequency of specific host-ARG pairs identified for most abundant bacterial genera in digester feed. Gammaproteobacterial genera *Stenotrophomonas* and *Acinetobacter* clustered separately and were characterized by species-specific multidrug resistance efflux complexes and the corresponding regulatory systems. *Stenotrophomonas* carried subunits for SmeABC and SmeDEF efflux pumps as well as the SmeRS regulatory system formerly identified in *S. maltophilia* (42, 43). *Acinetobacter* carried subunits for AdeIJK and AdeFGH multidrug efflux pumps together with multidrug resistance genes *adeB, abeM*, transcriptional activator *mexT* and aminoglycoside resistance gene *APH(3’)-Ia*. In addition to *Stenotrophomonas* and *Acinetobacter*, other identified ARG host genera in the digester feed included *Pseudomonas, Enterobacter, Klebsiella, Raoultella, Rahnella, Rouxiella, Pantoea, Serratia*, *Erwinia*, *Citrobacter*, *Leclercia* and *Lelliottia* that formed a network by sharing connections with ARGs belonging mainly to multidrug resistance type. For example, *Pseudomonas* was characterized by the highest number of ARG connections carrying subunits of the MexEF-OprN efflux system; similarly, *Enterobacter* and *Klebsiella* shared connections to subunits of MdtABC-TolC and AcrAB-TolC efflux systems. In addition to multidrug resistance genes, genes conferring resistance to polymyxin were related to multiple host genera in digester feed: *rosA* and *rosB* were both found in *Serratia, Rahnella* and *Rouxiella, arnA* was found in *Citrobacter* and *Pseudomonas, pmrE* in *Enterobacter* and *Klebsiella*, and *pmrA* in *Raoultella*.

**FIG 5.**
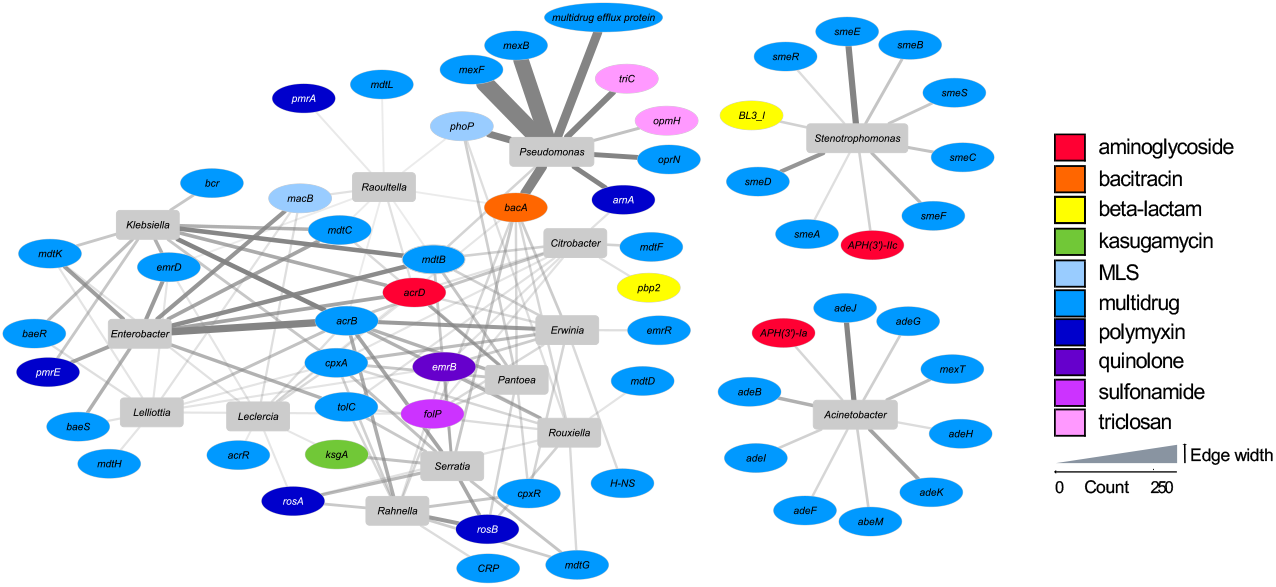
Most abundant bacterial host genera of non-plasmid ARGs detected in digester feed (FW1, FW2, LBF1, LBF2). ARGs were identified on MEGAHIT-assembled contigs by BLAST analysis of protein coding genes using local SARG reference database (Table S7 in supplemental material). Non-plasmid contigs carrying ARGs were further annotated for their taxonomic affiliation using Kaiju. ARGs are depicted in ellipses colored by resistance type and bacterial genera are depicted in grey rectangles. The width of the connecting edge corresponds to detection frequency of individual host-ARG pairs.

Only a limited number of ARGs were detected and their host organisms taxonomically determined in digestion products (Table 2). Bacterial hosts of ARGs in digestion products included genera formerly identified in anaerobic digesters associated with microbial degradation processes, such as *Fermentimonas*, and several sulphate-reducing genera *(Desulfobacca, Desulfococcus)*. More importantly, *Burkholderia* and *Arcobacter* (both carrying isoniazid resistance gene *katG*), *Streptococcus* (carrying MLS resistance gene *mefA*) and *Escherichia* (carrying MLS resistance gene *mefB*) were identified as ARG host organisms of clinical relevance.

**TABLE 2.** Bacterial host organisms of non-plasmid ARGs detected in digestion products (DG74, DG78, UASB). ARGs were identified on MEGAHIT-assembled contigs by BLAST analysis of protein coding genes using local SARG reference database (Table S7 in supplemental material). Non-plasmid contigs carrying ARGs were further annotated for their taxonomic affiliation using Kaiju.

| Kingdom/Phylum/Class | Family/Genus | ARG | ARG type |
| --- | --- | --- | --- |
| *Firmicutes* | not determined | *ANT(6)-Ia* | aminoglycoside |
| *Terrabacteria group* | not determined | *ANT(9)-Ia* | aminoglycoside |
| *Bacteroidetes* | *Proteiniphilum* | *bacA* | bacitracin |
| *Betaproteobacteria* | *Thauera* | *bacA* | bacitracin |
| *Deltaproteobacteria* | *Desulfomicrobium* | *bacA* | bacitracin |
| *Firmicutes* | *Alkaliphilus* | *bacA* | bacitracin |
| *Firmicutes* | *Thermoanaerobacterium* | *bcrA* | bacitracin |
| *Kiritimatiellaeota* | *Kiritimatiella* | *OXA-119* | beta-lactam |
| *Betaproteobacteria* | *Burkholderia* | *katG* | isoniazid |
| *Epsilonproteobacteria* | *Arcobacter* | *katG* | isoniazid |
| *Bacteroidetes* | *Fermentimonas* | *ermF* | MLS |
| *Firmicutes* | not determined | *linB* | MLS |
| *Deltaproteobacteria* | *Desulfococcus* | *lnuF* | MLS |
| *Deltaproteobacteria* | *Geobacter* | *lnuF* | MLS |
| *Firmicutes* | *Streptococcus* | *mefA* | MLS |
| *Gammaproteobacteria* | *Escherichia* | *mefB\** | MLS |
| *Firmicutes* | *Clostridium* | *mel* | MLS |
| *Firmicutes* | *Desulfitobacterium* | *vatB* | MLS |
| *Betaproteobacteria* | *Thauera* | *acrB* | multidrug |
| *Deltaproteobacteria* | *Desulfobacca* | *acrB* | multidrug |
| *Deltaproteobacteria* | *Syntrophobacter* | *acrB\** | multidrug |
| *Planctomycetes* | *Thermogutta* | *acrB* | multidrug |
| *Firmicutes* | not determined | *lsaE* | multidrug |
| *Deltaproteobacteria* | *Desulfobacca* | *smeE* | multidrug |
| *Gammaproteobacteria* | *Sedimenticola* | *smeR* | multidrug |
| *Betaproteobacteria* | *Roseateles* | *arnA* | polymyxin |
| *Gammaproteobacteria* | *Enterobacteriaceae* | *arnA* | polymyxin |
| *Bacteria* | not determined | *sulI* | sulfonamide |
| *Bacteria* | not determined | *sulI* | sulfonamide |

\* ARG was affiliated to the respective genus on several contigs

To further investigate the potential hosts of ARGs, the assembled contigs were binned into 201 metagenome assembled genomes (MAGs) (tools 3.11, 3.12 and 3.13 in Fig. 1) with >75% completeness and <25% redundancy (Table S9, Fig. S4 in supplemental material). Notably, MAGs found in digestion products (DG74, DG78, UASB) accounted for 75% of all assembled MAGs and did not contain ARGs. On the contrary, four MAGs containing ARGs were identified in digester feed (FW1, FW2, LBF1, LBF2) (Table 3). Interestingly, a genome belonging to the plant pathogenic genus *Erwinia* carried 19 ARGs conferring resistance to aminoglycoside, bacitracin, polymyxin, quinolone and sulfonamide antibiotics while also harboring several genes for multidrug resistance. Lactic acid bacteria *Lactococcus lactis* and *Lactobacillus* that are generally associated with probiotic features displayed resistance for MLS (*Lactococcus lactis* genes *lmrC, lmrD, lmrP*), tetracycline (*Lactococcus lactis* genes *tetM, tetS*) and trimethoprim (*Lactobacillus* gene *dfrE*). Additionally, a MAG belonging to the family *Bifidobacteriaceae* carried the inner membrane transporter gene *mdsB* of multidrug and metal efflux complex MdsABC. The four ARG-containing MAGs that were highly abundant in digester feed were not identified in digestion products (Fig. S4 in supplemental material).

**TABLE 3.**
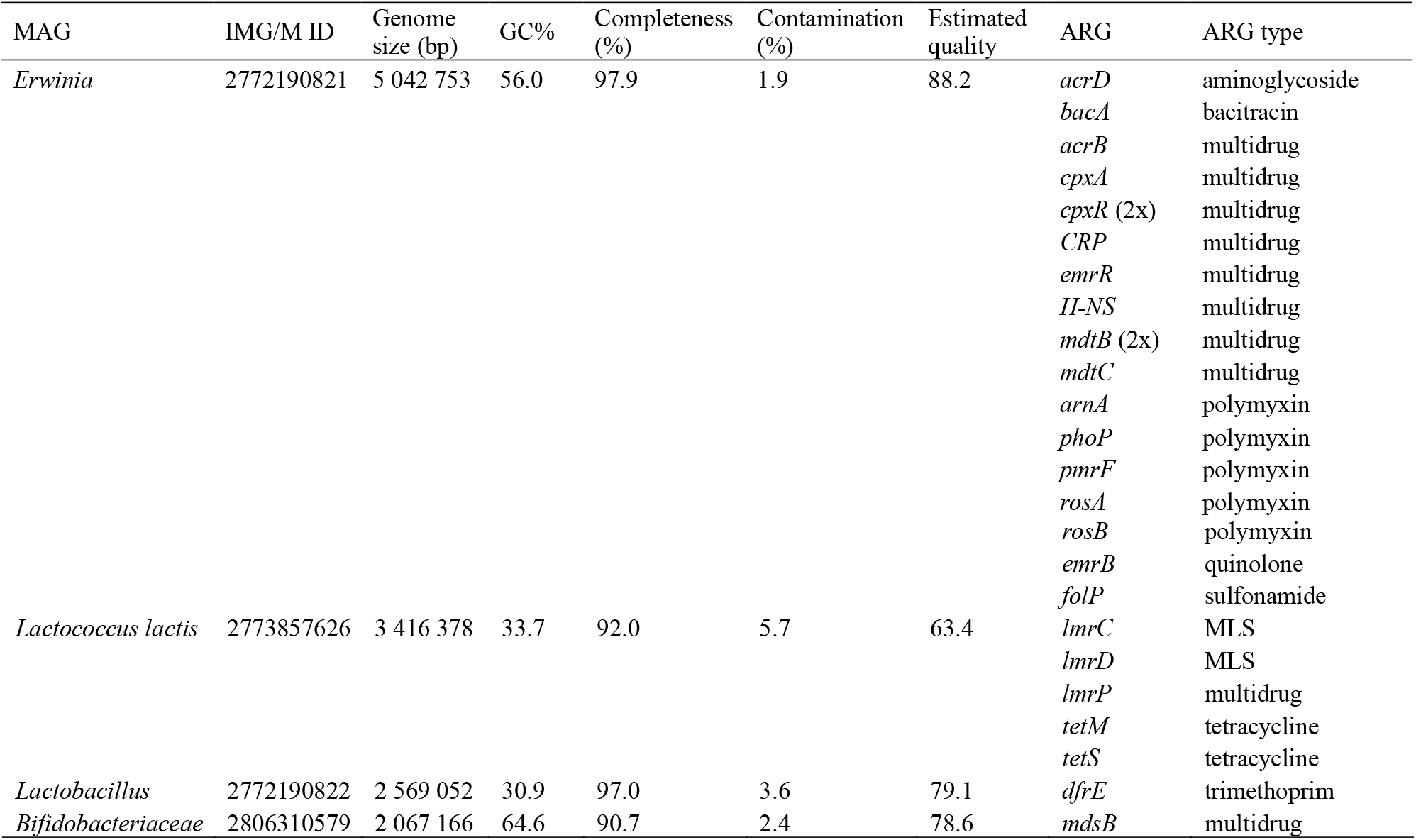
Metagenome assembled genomes (MAGs) containing antibiotic resistance genes (ARGs) originating from digester feed (FW1, FW2, LBF1, LBF2). ARGs were not detected in MAGs from digestion products. MAGs were produced by MaxBin and metaBAT, dereplicated with dRep and evaluated with CheckM for genome size, GC%, completeness, contamination and estimated quality. ARGs that were detected in two separate copies in a given MAG are indicated by parenthesis (2x). All ARG-containing MAGs have been deposited to the JGI Integrated Microbial Genomes and Microbiomes (IMG/M). Additional information about all MAGs is available in Table S9 and Figure S4 in supplemental material.

## DISCUSSION

AD is widely used for treatment of solid organic waste providing waste stabilization as well as energy (biogas) production. More recently, the potential of AD to reduce antibiotic resistant bacteria and ARGs has been investigated with mixed results (31). In this study, we examined the effect of co-digestion of food waste, paper and cardboard on microbial community composition and resistome using SSU rRNA gene sequencing, total community metagenome sequencing and quantitative PCR. The studied AD system that was a lab-scale solid-state leach bed reactor with leachate recycle via a UASB reactor that has been thoroughly described elsewhere (40). This AD system performed very well over 1.5 years of operation, also maintaining a stable microbial community (44).

### Microbial community composition of digester feed and digestion products

Microbial community composition has been suggested as one of the main drivers of ARGs in anaerobic digesters (35, 45). The microbial communities of the digester feed – a mix of food waste and lignocellulosic fibres – revealed aerobes and facultative aerobes that were distinct from the strictly anaerobic bacteria and archaea inside the digester. The FW alone and the blended LBF samples had very similar taxonomic profiles indicating that food waste was the main contributor to the microbial community in the digester feed, consisting mainly of microorganisms native to food products or of the microbes that colonized food waste during collection and storage prior to AD.

Previous studies on food microbiomes have highlighted the role of fermentative organisms such as lactic acid bacteria in fermentation processes (46). Several OTUs belonging to the genera *Lactobacillus* and *Leuconostoc* were identified in the food waste used in this study, indicating the presence of fermented food products (e.g. cheese, sourdough) or the start of degradation processes during the collection, storage and pre-processing of food waste prior to AD. *Lactobacillus* and *Leuconostoc* may also indicate spoilage of meat as previously shown for vacuum-packed pork (47), beef (48–50), minced meat (51) and sausages (52, 53). Besides lactic acid bacteria, several genera from the family *Enterobacteriaceae* were detected in digester feed, including *Citrobacter, Enterobacter, Erwinia, Kluyvera, Pantoea, Serratia* and *Lelliottia*. Jackson *et al*. (54) identified *Serratia*, *Erwinia*, *Enterobacter* and *Pantoea* from leafy salad vegetables, which are also common in food waste. While many of the identified *Enterobacteriaceae* are widely recognized as plant pathogens or symbionts (e.g. *Erwinia*), potential human pathogens were also detected (e.g. *Enterobacter, Serratia*).

Special attention should be given to the occurrence of ESKAPE pathogens – *Enterococcus faecium, Staphylococcus aureus, Klebsiella pneumoniae, Acinetobacter baumannii, Pseudomonas aeruginosa*, and *Enterobacter* species, that are common causes of nosocomial infections and characterized by various antimicrobial resistance mechanisms (55). A total of 15 OTUs annotated to *S. aureus, K. pneumoniae, A. baumannii* or *P. aeruginosa* were found in very low abundance in digester feed (each with relative abundance below 0.12% of total OTUs), while none of these were detected in digestion products. One hundred and two (102) OTUs were annotated to *Enterobacter* comprising up to 0.13% of OTUs in digester feed, while none was detected in digestion products. Thus, ESKAPE pathogens were detectable in relatively low abundance in the digester feed used in this study but did not survive anaerobic treatment.

The microbial community after AD was completely different from that in the feed. Hydrolytic, fermenting, acidogenic, acetogenic and methanogenic microorganisms comprise a typical microbial community in a stable AD process. Representatives from the phyla *Actinobacteria*, *Bacteroidetes*, *Chloroflexi*, *Firmicutes* and *Proteobacteria* that have been connected to hydrolysis, fermentation and acidogenesis (56) were identified in the bacterial communities of the digestate samples in this study. As expected, digestates exhibited abundant archaeal communities responsible for methane production. Acetoclastic methanogen *Methanosaeta* was identified as the dominant archaeal genus (reaching up to 72.6% of total archaeal OTUs in DG74), which has been noted before for digesters with high acetate concentration (57, 58). Members of the orders *Methanobacteriales* and *Methanomicrobiales* commonly identified in anaerobic digesters treating food waste (59–61) were also present in the digestates of the current study. Thus, a diverse methanogenic community achieved stable methane production, as evidenced by substantial feedstock degradation and biogas production rates (40).

### ARGs in digester feed and digestion products

ARGs were identified in all samples, in digester feed as well as in AD products. Higher richness as well as higher relative abundance of individual ARGs per 16S rRNA gene was found in digester feed compared to digestion products indicating the ability of AD to reduce ARG diversity and abundance. Total relative abundance of ARGs per 16S rRNA gene reached 0.84 ARG/16S rRNA gene in digester feed (LBF1), which is comparable to ARG levels in faeces and wastewater from livestock farms (36), while the respective value in digestion products was below 0.05 ARG/16S rRNA (0.04 ARG/16S rRNA in UASB), comparable to natural environments such as soils, sediments and river water (36). Thus, AD clearly reduced ARG values from levels associated with high contamination to environmental background levels.

Yet, although AD significantly reduced the abundance of ARGs, a limited number of ARGs were also detected in digestion products. Total community metagenome sequencing and HT-qPCR array both detected ARGs in digestion products that were low in abundance relative to the total microbial population (as shown by total community metagenome sequencing), but achieved high absolute abundance per gram dry mass of digestate (reaching 10^8^ copies/g-dw for individual genes and 10^9^ copies/g-dw in total) due to high bacterial loads in the digestates (as shown by HT-qPCR array). Same order of magnitude absolute abundance values of individual ARGs detected by qPCR have also been reported previously for anaerobic digestates of FW (62). Thus, the effectiveness of AD on treating waste containing ARGs is a function of which microbes can grow in the AD system. In this case study, ARGs from food waste microbes were eliminated while ARGs present in the microbes involved in the digestion process were more abundant in the final digestate, and were at high absolute abundance owing to the high concentration of microbes generally after digestion. Future experiments should focus on post-treatment of digested solids, such as additional aerobic curing to reduce odor and stabilize waste, to determine the ultimate environmental load of ARGs associated with land application of anaerobic digestates.

Special attention should be given to the mobility of ARGs that can be transferred from environmental bacteria to human pathogens or vice versa as shown previously (63). In this study, on average, 32% of ARGs in assembled contigs from digester feed were located on plasmids, indicating their potential mobility. Similar plasmid proportions from total assembled contigs have also been found in photobioreactor microbial communities (64) and microbial mats inhabiting mine waters (65). The HT-qPCR array detected a wide array of MGEs including insertional sequences, integrases, transposases and plasmids mostly in food waste but also in digestates. Class 1 integron-integrase gene *intI1* has been suggested as a proxy for anthropogenic pollution (66) and was also found in digester feed (in the range of 10^5^–10^6^ copies/g-dw) as well as in digestion products (10^8^ copies/g-dw) of this study. The occurrence of *intI1* in digestion products correlates with the limited number of ARGs detected in the digestates of this study, showing that anthropogenic impact remains detectable after AD.

### Potential bacterial host organisms of ARGs

Correlation and network analysis have been used in previous studies to link ARGs to their potential host organisms (35–37, 67). Here we used a combination of metagenomic assembly and binning methods to detect taxonomically annotated contigs and MAGs with ARGs.

Taxonomic annotation of ARG-containing contigs showed several connections between bacterial genera and ARGs. Among others, the genera *Enterobacter, Klebsiella, Acinetobacter* and *Pseudomonas* that include medically critical ESKAPE-pathogens were connected to ARGs in digester feed conferring resistance to multidrug, MLS, polymyxin, aminoglycoside, triclosan and bacitracin antibiotics. The high number of multidrug resistance genes is especially worrisome as these genes have the potential to confer resistance to several types of antibiotics. Besides that, 12 out of 14 of the most abundant ARG-containing genera in digester feed also included resistance genes to polymyxin, a last-resort antibiotic used against Gram-negative bacteria. This highlights the spread of ARGs against last-resort antibiotics among various bacterial genera. ARG-containing genera that include clinically important species were also detected in digestion products, including *Burkholderia, Arcobacter, Streptococcus* and *Escherichia* indicating the risks associated with the possible use of anaerobic digestates.

Using metagenomic binning we were able to detect four ARG-containing MAGs originating from digester feed. A MAG annotated as *Erwinia* was found to carry 19 ARGs conferring resistance to several classes of antibiotics. Although *Erwinia* is commonly known as a plant pathogen, few cases of human infections have also been reported (68, 69). As *Erwinia* is closely related to other *Enterobacteriaceae* that include several known human pathogens, horizontal transfer of ARGs from plant-pathogenic *Erwinia* to human pathogenic genera may occur. Additionally, MAGs annotated as lactic acid bacteria *Lactococcus lactis* and *Lactobacillus* were found to carry ARGs. These bacteria are extensively used for food fermentation processes and are often regarded as probiotics, however, increasing evidence suggests they are reservoirs of potentially transmissible ARGs and may play a crucial role in the acquisition of AMR via food (70). It is noteworthy, that although 75% of all MAGs originated from digestion products, these MAGs did not contain ARGs. This correlates with the low abundance of ARGs in digestion products detected by other methods used in this study.

### Methodological aspects of ARG detection

In this study total community metagenome data and HT-qPCR data were used to characterize the diversity and abundance of ARGs. While the metagenomic approach has the potential to detect higher diversity of ARGs due to large reference databases, qPCR of ARGs, although being limited in the number of ARGs even in high-throughput applications, can provide higher sensitivity (71). Our results showed that both metagenomic and HT-qPCR approach detected a broad spectrum of ARGs with higher diversity of ARGs in digester feed compared to digestion products. Among the limited number of ARGs detected in digestion products *mphA, lnuB, catQ* and *tet44* were found to be unique to digestate by both metagenomic and HT-qPCR approach. Additionally, HT-qPCR provided absolute abundance values for individual ARGs that could be related back to the number of gene copies present per gram of digester feed and in digested solids. This type of quantitative information on ARGs is required for assessing the risks associated with food waste and the use of anaerobic digestates as agricultural fertilizers. Thus, a combination of shotgun metagenomic and qPCR approach is recommended for a comprehensive view of a sample’s resistome and for the assessment of AMR risks.

In addition to the annotation of short sequencing reads, we used assembled contigs to provide genomic context of ARGs. The distribution of ARG types on assembled contigs closely resembled that on short sequencing reads, additionally, assembled data provided information about genomic location and potential host organisms of ARGs, which cannot be determined from short sequencing reads. Although assembled contigs provide better resolution for taxonomic and functional annotation than short sequencing reads, assembly may also introduce biases (71). Similar problem has been noted for metagenomic binning, which often has limited power for analysis of complex microbial communities. This was also evident in the results of the current study with 75% of MAGs (151 MAGs) originating from digestion products and only 25% of MAGs (50 MAGs) originating from the heterogeneous microbial communities of digester feed. Thus, our findings should be interpreted in light of the limitations of current methods used for taxonomic and functional analysis of complex microbial communities.

In conclusion, using a combination of metagenome sequencing, assembly and binning as well as quantitative PCR analysis is recommended to estimate the diversity and abundance of ARGs and to situate the ARGs within their genomic context and potential host organisms. The detection of potential host organisms and genomic context allows for a more accurate risk evaluation associated with solid organic wastes.

## MATERIALS AND METHODS

### Anaerobic digestion system and feedstock

A lab-scale anaerobic digestion system designed for the treatment of solid organic waste was operated for a total of 88 weeks. System design, its operating parameters and feedstock have been described in detail by Guilford *et al*. (40). In short, the anaerobic digestion system consisted of 6 leach beds (8.5 L each), an upflow anaerobic sludge blanket reactor (UASB, 27.5 L) treating leach bed leachate, a UASB feed tank (17.5 L), a leach bed feed tank (17.5 L), three peristaltic pumps to recirculate leachate, two wet-tip gas meters for biogas measurement, and an automated control system. The system was maintained at 37–39°C with continuous recirculation of the leachate. It was operated in sequential batch feeding mode: each week one of the leach beds was filled with a mixture of lignocellulosic fibres and food waste (FW) making up the leach bed feed (LBF), recovered from local residential waste recycling programs. Lignocellulosic fibres included shredded cardboard, boxboard, fine paper and newsprint; food waste was collected from a residential green bin program and sorted manually to remove bones and inorganic materials. Solids retention time in the system was six weeks.

### Sampling and DNA extraction

Samples for microbial community analysis were collected during weeks 75–84 of the experiment, during which FW contributed 21.3% of the total COD in the feedstock and the system exhibited stable performance with a methane yield of 246 L-CH4 kg-VS_added_^-1^ and a substrate destruction efficiency, as volatile solids (VS), of 63.5% (40). Ten 50 mL samples were collected: three food waste samples FW1, FW2, FW3 (FW source and composition has been described by Guilford *et al*. (40)); three leach bed feed samples LBF1, LBF2, LBF3 consisting of the respective food waste mixed with lignocellulosic fibres; three digestate samples DG74, DG76, DG78 collected after 6 weeks of digestion; and one sample from the microbial granular sludge of the UASB reactor (UASB). It should be noted that DG74, DG76 and DG78 were not direct digestion products of LBF1, LBF2 and LBF3, respectively, although derived from the same source of FW and lignocellulosic fibres used for LBF1, LBF2 and LBF3. All samples were preserved at –20°C until DNA extraction.

Total community DNA was extracted using the PowerMax Soil DNA Isolation Kit (MoBio Laboratories, Carlsbad, CA, USA) from 5 g of sample according to manufacturer’s protocol. FW and LBF samples were further purified with 5 M NaCl and 100% ethanol as recommended by the manufacturer. The quantity and quality of DNA extracts was confirmed using NanoDrop spectrophotometer ND-1000 (Wilmington, DE, USA). All DNA extracts were stored at –80°C.

### Quantitative PCR analyses

Quantitative PCR (qPCR) was used to quantify total bacterial (method 1.1 in Fig. 1) and archaeal (method 1.2 in Fig. 1) 16S rRNA gene copies using a CFX96 Touch Real-Time PCR Detection System (Bio-Rad Laboratories, Hercules, CA, USA) and previously published primers targeting most bacterial 16S rRNA genes: Bac1055f (5’- ATGGCTGTCGTCAGCT-3’) and Bac1392r (5’- ACGGGCGGTGTGTAC-3’) (72, 73), and most archaeal 16S rRNA genes: Arch787f (5’-ATTAGATACCCGBGTAGTCC-3’) and Arch1059r (5’-GCCATGCACCWCCTCT-3’) (74). qPCR reactions were performed in 20 μL comprising 10 μL SsoFast EvaGreen Supermix (Bio-Rad Laboratories, Hercules, CA, USA), 0.5 μM each forward and reverse primer, 2 μL template DNA, and sterile UltraPure distilled water. The thermocycling program was as follows: initial denaturation at 98°C for 2 min, followed by 40 cycles of denaturation at 98°C for 5 s, annealing at Tm (55°C for bacterial 16S rRNA, 60°C for archaeal 16S rRNA) for 10 s, and final melting curve analysis in the range of 65–95°C (steps of 0.5°C for 5 s). All samples were measured in three technical replicates and every qPCR run included negative controls containing all reaction components except for template DNA. Serial dilutions of plasmid stocks containing corresponding 16S rRNA gene fragments were used to generate standard curves. Amplification data was analyzed using CFX Manager Software v.3.1. Copy numbers of bacterial and archaeal 16S rRNA genes per gram of sample dry weight were calculated as follows: Absolute Abundance = StartingQuantity * DNAElutionVolume / (SampleWeight*SampleDW) where StartingQuantity represents the gene copies in 1 μL DNA extract as estimated by the CFX Manager Software, DNAElutionVolume was 5000 μL for DG74, DG76, DG78, UASB and 100 μL for FW1, FW2, FW3, LBF1, LBF2, LBF3, SampleWeight was the mass of the sample used for DNA extraction and SampleDW was the dry weight of the sample material used for DNA extraction.

A high-throughput qPCR (HT-qPCR) array was used for the quantification of ARGs (method 4.1 in Fig. 1) and MGEs (method 4.2 in Fig. 1) in FW1, FW2, DG74 and DG78. DNA extracts were sent to Dr Robert Stedtfeld at Michigan State University (USA) who performed the analysis. A total of 384 primer sets were used targeting 315 ARGs and 57 MGEs, additionally taxonomic marker genes were included in the array (75). All HT-qPCR reactions were performed using Takara (previously WaferGen) SmartChip real-time PCR system as described previously (76). In brief, 5184 parallel qPCR reactions (100 nL) were dispensed into the SmartChip using the SmartChip Multisample Nanodispenser, followed by thermal cycling in the SmartChip Cycler, automatic melting process and initial data processing with SmartChip qPCR Software (v.2.7.0.1) as described before (76). Wells with multiple melt peaks and wells with amplification efficiency outside the range 1.75 to 2.25 were removed from analysis. Each sample was analyzed by three technical replicates. Genes detected in only one of the replicates were considered false positives and removed from analysis. Genomic copy numbers were estimated using the formula in Looft *et al*. (77) with the exception of setting detection limit to threshold cycle (Ct) 28 as suggested by Stedtfeld *et al*. (75). Relative abundances of the studied genes were calculated as ratios to 16S rRNA: RelativeAbundance_gene_ = GenomicCopyNumber_gene_ / GenomicCopyNumber_16S_. Absolute abundances were determined by multiplying the relative abundance of a gene by 16S rRNA absolute abundance (determined by regular qPCR analysis described above): AbsoluteAbundance_gene_ = RelativeAbundance_gene_ * AbsoluteAbundance_16S_. ARG absolute abundances were visualized in R software using the package ComplexHeatmap (78).

### SSU rRNA gene amplicon sequencing and taxonomic analyses

SSU rRNA gene sequencing with the universal primers 926f-modified (5’-AAACTYAAAKGAATWGRCGG-3’) and 1392r-modified (5’-ACGGGCGGTGWGTRC-3’) (modified from (79)) targeting the V6–V8 variable region of the 16S rRNA gene from bacteria and archaea as well as the 18S rRNA gene in eukarya was performed at McGill University and Génome Québec Innovation Centre (Canada) on Illumina MiSeq System (PE300). The analysis of sequencing reads was performed with QIIME1 package (tool 2.1 in Fig. 1) (80) by joining forward and reverse reads (multiple_join_paired_ends.py, min overlap 50 bp, max mismatch allowance 8), removing low-quality sequences (multiple_split_libraries_fastq.py, quality threshold 19), removing chimeric sequences and identifying operational taxonomic units at 97% similarity (identify_chimeric_seqs.py, pick_open_reference_otus.py, method usearch61 (81), reference database SILVA release 128).

### Total community metagenome sequencing and bioinformatic workflow

Total community metagenome sequencing of the samples FW1, FW2, LBF1, LBF2, DG74, DG78, UASB was performed at University of Toronto Centre for the Analysis of Genome Evolution and Function (Canada) on Illumina NextSeq500 Desktop Sequencer v2 using High Output flowcell (PE150). Bioinformatic workflow included quality control and trimming of the sequencing reads, taxonomic classification, metagenomic assembly and binning, and detection of ARGs. Sequence reads were trimmed with Trimmomatic v.0.32 (82) (tool 3.1 in Fig. 1) using default settings for paired-end mode, additional trimming to remove polyG sequences (≥30 bp) was performed with AfterQC v.0.9.1 (83) (tool 3.2 in Fig. 1).

Quality-trimmed reads were subjected to taxonomic classification using Kaiju v.1.4.5 (84) (tool 3.3 in Fig. 1) in greedy mode (allowed substitutions 5, minimum required match length 11, minimum required match score 70) with bacterial, archaeal, eukaryotic and viral protein sequences from the NCBI nr database as reference. To further investigate microbial community composition, metaxa2 v.2.1 (85) (tool 3.4 in Fig. 1) was used to extract small subunit rRNA gene reads in metagenome mode with default search criteria and reliability score cut-off 80.

Quality-trimmed reads were assembled into longer DNA contigs using MEGAHIT v.1.1.1 (86) (tool 3.6 in Fig. 1) and metaSPAdes v.3.10.1 (87) (tool 3.7 in Fig. 1) with default settings. Assemblies were compared using metaQUAST v.4.5 (88) which also provided assembly statistics. Metagenome binning was performed using MaxBin v.2.2.3 (89) (tool 3.11 in Fig. 1) and MetaBAT v.2.12 (90) (tool 3.12 in Fig. 1) with default settings on the MEGAHIT assemblies. All genome bins produced by both binning methods were dereplicated using dRep v.1.4.3 (91) (tool 3.13 in Fig. 1) with default settings (ANI cut-off 99%, minimum genome completeness 75%, maximum contamination 25%) and evaluated with CheckM v.1.0.7 (92). Quality of the resulting metagenome-assembled genomes (MAGs) was defined as (Completeness – 5*Contamination) as suggested by Parks *et al*. (93). MAGs with quality >50 and taxonomic classification to at least phylum level, were submitted to the JGI Integrated Microbial Genomes and Microbiomes (IMG) database for annotation. Anvi’o v.5 (94) was used for visualizing MAGs clustered based on the relative abundance of MAGs in the samples: Euclidian distances were calculated from relative abundance estimates of each MAG and clustering was performed using the ward linkage algorithm. Relative abundance of each MAG was calculated as number of reads recruited to the MAG divided by total reads recruited to that MAG across all samples.

ARGs from subsets of quality-trimmed sequencing reads (20 million reads/sample) were determined with ARGs-OAP v.1.2 (41) (tool 3.5 in Fig. 1) using the default reference database SARG and default settings. The same reference database was also used to detect ARGs on assembled contigs by local BLASTp (evalue 1e-7, identity percent 80, minimal alignment length 25) using protein coding genes annotated by JGI IMG/M 4 version (95) as query sequences (tool 3.10 in Fig. 1). Contigs with multiple ARGs were identified by a local python script that counted how many times a contig ID appeared in the list of contigs that carried ARGs (Method S1 in supplemental material). PlasFlow v.1.1 (65) with default settings was used to predict plasmid sequences from the contigs with ARGs (tool 3.8 in Fig. 1). Taxonomic classification of the non-plasmid contigs carrying ARGs was determined with Kaiju v.1.4.5 (84) in greedy mode (maximum mismatches allowed 5, minimal required match length 11, minimal required match score 70, filtering of query sequences containing low-complexity regions on) (tool 3.9 in Fig. 1) using NCBI RefSeq reference database and visualized with Cytoscape v.3.6.1 (96) by selecting 10 most abundant ARGs from all genera above relative abundance of 1%. The presence of ARGs in MAGs was determined by a local python script comparing a list of ARG-carrying contigs to the list of contigs included in the MAG (tool 3.14 in Fig. 1, Method S2 in supplemental material).

### Accession numbers

Sequencing data from SSU rRNA gene sequencing (accession numbers provided in Table S1 in supplemental material) and total community metagenome sequencing (accession numbers provided in Table S3 in supplemental material) is available on NCBI Sequence Read Archive under BioProject PRJNA501900, SRA study SRP167436, accession numbers SRX4965138–47 (SSU rRNA gene sequencing) and SRX4986160–6 (total community metagenome sequencing). Additionally, metagenome assemblies (accession numbers provided in Table S6 in supplemental material) and metagenome assembled genomes (accession numbers provided in Table S9 in supplemental material) have been deposited to the JGI Integrated Microbial Genomes and Microbiomes (https://img.jgi.doe.gov/cgi-bin/m/main.cgi).

## Supporting information

Method S1_python script

Method S2_python script

## ACKNOWLEDGEMENTS

This research was funded by the Natural Sciences and Engineering Research Council of Canada (Collaborative Research and Development Grant), and by Miller Waste Systems Inc. We would like to thank Dr. Robert Stedtfeld from Michigan State University for his help advising and running the high-throughput qPCR analysis.

## REFERENCES

1. Pepper IL, Brooks JP, Gerba CP. 2018. Antibiotic Resistant Bacteria in Municipal Wastes: Is There Reason for Concern? Environ Sci Technol 52:3949–3959.

2. WHO. 2015. Global action plan on antimicrobial resistance. World Health Organization, Geneva, Switzerland.

3. O’Neill J. 2016. Tackling Drug-Resistant Infections Globally: Final Report and Recommendations. The Review on Antimicrobial Resistance. HM Government and Wellcome Trust.

4. Robinson TP, Bu DP, Carrique-Mas J, Fèvre EM, Gilbert M, Grace D, Hay SI, Jiwakanon J, Kakkar M, Kariuki S, Laxminarayan R, Lubroth J, Magnusson U, Thi Ngoc P, Van Boeckel TP, Woolhouse MEJ. 2016. Antibiotic resistance is the quintessential One Health issue. Trans R Soc Trop Med Hyg 110:377–80.

5. Bengtsson-Palme J, Kristiansson E, Larsson DGJ. 2018. Environmental factors influencing the development and spread of antibiotic resistance. FEMS Microbiol Rev 42.

6. Huijbers PMC, Blaak H, de Jong MCM, Graat EAM, Vandenbroucke-Grauls CMJE, de Roda Husman AM. 2015. Role of the Environment in the Transmission of Antimicrobial Resistance to Humans: A Review. Environ Sci Technol 49:11993–12004.

7. D’Costa VM, King CE, Kalan L, Morar M, Sung WWL, Schwarz C, Froese D, Zazula G, Calmels F, Debruyne R, Golding GB, Poinar HN, Wright GD. 2011. Antibiotic resistance is ancient. Nature 477:457–61.

8. Durso LM, Wedin DA, Gilley JE, Miller DN, Marx DB. 2016. Assessment of Selected Antibiotic Resistances in Ungrazed Native Nebraska Prairie Soils. J Environ Qual 45:454–462.

9. Perron GG, Whyte L, Turnbaugh PJ, Goordial J, Hanage WP, Dantas G, Desai MM. 2015. Functional Characterization of Bacteria Isolated from Ancient Arctic Soil Exposes Diverse Resistance Mechanisms to Modern Antibiotics. PLoS One 10:e0069533.

10. Hatosy SM, Martiny AC. 2015. The ocean as a global reservoir of antibiotic resistance genes. Appl Environ Microbiol 81:7593–9.

11. Czekalski N, Sigdel R, Birtel J, Matthews B, Bürgmann H. 2015. Does human activity impact the natural antibiotic resistance background? Abundance of antibiotic resistance genes in 21 Swiss lakes. Environ Int 81:45–55.

12. Berendonk TU, Manaia CM, Merlin C, Fatta-Kassinos D, Cytryn E, Walsh F, Bürgmann H, Sørum H, Norström M, Pons M-N, Kreuzinger N, Huovinen P, Stefani S, Schwartz T, Kisand V, Baquero F, Martinez JL. 2015. Tackling antibiotic resistance: the environmental framework. Nat Rev Microbiol 13:310–7.

13. Braguglia CM, Gallipoli A, Gianico A, Pagliaccia P. 2018. Anaerobic bioconversion of food waste into energy: A critical review. Bioresour Technol 248:37–56.

14. Hoornweg D, Bhada-Tata P. 2012. What a Waste : A Global Review of Solid Waste Management. World Bank, Washington, DC.

15. Friedman M. 2015. Antibiotic-Resistant Bacteria: Prevalence in Food and Inactivation by Food-Compatible Compounds and Plant Extracts. J Agric Food Chem 63:3805–3822.

16. Bengtsson-Palme J. 2017. Antibiotic resistance in the food supply chain: where can sequencing and metagenomics aid risk assessment? Curr Opin Food Sci 14:66–71.

17. Chajęcka-Wierzchowska W, Zadernowska A, Łaniewska-Trokenheim Ł. 2016. Diversity of Antibiotic Resistance Genes in Enterococcus Strains Isolated from Ready-to-Eat Meat Products. J Food Sci 81:M2799–M2807.

18. Noyes NR, Yang X, Linke LM, Magnuson RJ, Dettenwanger A, Cook S, Geornaras I, Woerner DE, Gow SP, McAllister TA, Yang H, Ruiz J, Jones KL, Boucher CA, Morley PS, Belk KE. 2016. Resistome diversity in cattle and the environment decreases during beef production. Elife 5:e13195.

19. Yang X, Noyes NR, Doster E, Martin JN, Linke LM, Magnuson RJ, Yang H, Geornaras I, Woerner DR, Jones KL, Ruiz J, Boucher C, Morley PS, Belk KE. 2016. Use of Metagenomic Shotgun Sequencing Technology To Detect Foodborne Pathogens within the Microbiome of the Beef Production Chain. Appl Environ Microbiol 82:2433–43.

20. Oliveira M, Viñas I, Usall J, Anguera M, Abadias M. 2012. Presence and survival of Escherichia coli O157:H7 on lettuce leaves and in soil treated with contaminated compost and irrigation water. Int J Food Microbiol 156:133–140.

21. Rahube TO, Marti R, Scott A, Tien Y-C, Murray R, Sabourin L, Zhang Y, Duenk P, Lapen DR, Topp E. 2014. Impact of fertilizing with raw or anaerobically digested sewage sludge on the abundance of antibiotic-resistant coliforms, antibiotic resistance genes, and pathogenic bacteria in soil and on vegetables at harvest. Appl Environ Microbiol 80:6898–907.

22. Holvoet K, Sampers I, Callens B, Dewulf J, Uyttendaele M. 2013. Moderate prevalence of antimicrobial resistance in Escherichia coli isolates from lettuce, irrigation water, and soil. Appl Environ Microbiol 79:6677–83.

23. Marti R, Scott A, Tien Y-C, Murray R, Sabourin L, Zhang Y, Topp E. 2013. Impact of Manure Fertilization on the Abundance of Antibiotic-Resistant Bacteria and Frequency of Detection of Antibiotic Resistance Genes in Soil and on Vegetables at Harvest. Appl Environ Microbiol 79:5701–5709.

24. Diarra MS, Delaquis P, Rempel H, Bach S, Harlton C, Aslam M, Pritchard J, Topp E. 2014. Antibiotic Resistance and Diversity of Salmonella enterica Serovars Associated with Broiler Chickens. J Food Prot 77:40–49.

25. Yang B, Cui Y, Shi C, Wang J, Xia X, Xi M, Wang X, Meng J, Alali WQ, Walls I, Doyle MP. 2014. Counts, Serotypes, and Antimicrobial Resistance of Salmonella Isolates on Retail Raw Poultry in the People’s Republic of China. J Food Prot 77:894–902.

26. Elhadi N. 2014. Prevalence and antimicrobial resistance of Salmonella spp. in raw retail frozen imported freshwater fish to Eastern Province of Saudi Arabia. Asian Pac J Trop Biomed 4:234–238.

27. Moore JE, Huang J, Yu P, Ma C, Moore PJ, Millar BC, Goldsmith CE, Xu J. 2014. High diversity of bacterial pathogens and antibiotic resistance in salmonid fish farm pond water as determined by molecular identification employing 16S rDNA PCR, gene sequencing and total antibiotic susceptibility techniques. Ecotoxicol Environ Saf 108:281–286.

28. Silveira-Filho VM, Luz IS, Campos APF, Silva WM, Barros MPS, Medeiros ES, Freitas MFL, Mota RA, Sena MJ, Leal-Balbino TC. 2014. Antibiotic Resistance and Molecular Analysis of Staphylococcus aureus Isolated from Cow’s Milk and Dairy Products in Northeast Brazil. J Food Prot 77:583–591.

29. Lee J, Shin SG, Jang HM, Kim YB, Lee J, Kim YM. 2017. Characterization of antibiotic resistance genes in representative organic solid wastes: Food waste-recycling wastewater, manure, and sewage sludge. Sci Total Environ 579:1692–1698.

30. Vasco-Correa J, Khanal S, Manandhar A, Shah A. 2018. Anaerobic digestion for bioenergy production: Global status, environmental and techno-economic implications, and government policies. Bioresour Technol 247:1015–1026.

31. Youngquist CP, Mitchell SM, Cogger CG. 2016. Fate of Antibiotics and Antibiotic Resistance during Digestion and Composting: A Review. J Environ Qual 45:537.

32. Pu C, Liu H, Ding G, Sun Y, Yu X, Chen J, Ren J, Gong X. 2018. Impact of direct application of biogas slurry and residue in fields: In situ analysis of antibiotic resistance genes from pig manure to fields. J Hazard Mater 344:441–449.

33. Zhang J, Mao F, Loh K-C, Gin KY-H, Dai Y, Tong YW. 2018. Evaluating the effects of activated carbon on methane generation and the fate of antibiotic resistant genes and class I integrons during anaerobic digestion of solid organic wastes. Bioresour Technol 249:729–736.

34. Zhang J, Chen M, Sui Q, Wang R, Tong J, Wei Y. 2016. Fate of antibiotic resistance genes and its drivers during anaerobic co-digestion of food waste and sewage sludge based on microwave pretreatment. Bioresour Technol 217:28–36.

35. Luo G, Li B, Li L-G, Zhang T, Angelidaki I. 2017. Antibiotic Resistance Genes and Correlations with Microbial Community and Metal Resistance Genes in Full-Scale Biogas Reactors As Revealed by Metagenomic Analysis. Environ Sci Technol 51:4069–4080.

36. Li B, Yang Y, Ma L, Ju F, Guo F, Tiedje JM, Zhang T. 2015. Metagenomic and network analysis reveal wide distribution and co-occurrence of environmental antibiotic resistance genes. ISME J 9:2490–2502.

37. Jang HM, Shin J, Choi S, Shin SG, Park KY, Cho J, Kim YM. 2017. Fate of antibiotic resistance genes in mesophilic and thermophilic anaerobic digestion of chemically enhanced primary treatment (CEPT) sludge. Bioresour Technol 244:433–444.

38. Stokes HW, Gillings MR. 2011. Gene flow, mobile genetic elements and the recruitment of antibiotic resistance genes into Gram-negative pathogens. FEMS Microbiol Rev 35:790–819.

39. Guilford NGH. 2017. The Anaerobic Digestion of Organic Solid Wastes of Variable Composition. PhD Thesis. University of Toronto.

40. Guilford NGH, Lee HP, Kanger K, Meyer T, Edwards EA. 2019. Solid state anaerobic digestion of mixed organic waste: the synergistic effect of food waste addition on the destruction of paper and cardboard. bioRxiv https://doi.org/10.1101/564203.

41. Yang Y, Jiang X, Cai B, Ma L, Li B, Zhang A, Cole JR, Tiedje JM, Zhang T. 2016. ARGs-OAP: Online Analysis Pipeline for Antibiotic Resistance Genes Detection from Metagenomic Data Using an Integrated Structured ARG-database. Bioinformatics 32:2346–2351.

42. Alonso A, Martínez JL. 2000. Cloning and characterization of SmeDEF, a novel multidrug efflux pump from Stenotrophomonas maltophilia. Antimicrob Agents Chemother 44:3079–86.

43. Li X-Z, Zhang L, Poole K. 2002. SmeC, an outer membrane multidrug efflux protein of Stenotrophomonas maltophilia. Antimicrob Agents Chemother 46:333–43.

44. Lee H. 2018. Characterization of the Microbial Community in a Sequentially Fed Anaerobic Digester Treating Solid Organic Waste. University of Toronto.

45. Miller JH, Novak JT, Knocke WR, Young K, Hong Y, Vikesland PJ, Hull MS, Pruden A. 2013. Effect of Silver Nanoparticles and Antibiotics on Antibiotic Resistance Genes in Anaerobic Digestion. Water Environ Res 85:411–421.

46. De Filippis F, Parente E, Ercolini D. 2018. Recent Past, Present, and Future of the Food Microbiome. Annu Rev Food Sci Technol 9:589–608.

47. Nieminen TT, Dalgaard P, Björkroth J. 2016. Volatile organic compounds and Photobacterium phosphoreum associated with spoilage of modified-atmosphere-packaged raw pork. Int J Food Microbiol 218:86–95.

48. Jääskeläinen E, Hultman J, Parshintsev J, Riekkola M-L, Björkroth J. 2016. Development of spoilage bacterial community and volatile compounds in chilled beef under vacuum or high oxygen atmospheres. Int J Food Microbiol 223:25–32.

49. Säde E, Penttinen K, Björkroth J, Hultman J. 2017. Exploring lot-to-lot variation in spoilage bacterial communities on commercial modified atmosphere packaged beef. Food Microbiol 62:147–152.

50. Ferrocino I, Greppi A, La Storia A, Rantsiou K, Ercolini D, Cocolin L. 2016. Impact of Nisin-Activated Packaging on Microbiota of Beef Burgers during Storage. Appl Environ Microbiol 82:549–59.

51. Stoops J, Ruyters S, Busschaert P, Spaepen R, Verreth C, Claes J, Lievens B, Van Campenhout L. 2015. Bacterial community dynamics during cold storage of minced meat packaged under modified atmosphere and supplemented with different preservatives. Food Microbiol 48:192–199.

52. Fougy L, Desmonts M-H, Coeuret G, Fassel C, Hamon E, Hézard B, Champomier-Vergès M-C, Chaillou S. 2016. Reducing Salt in Raw Pork Sausages Increases Spoilage and Correlates with Reduced Bacterial Diversity. Appl Environ Microbiol 82:3928–3939.

53. Benson AK, David JRD, Gilbreth SE, Smith G, Nietfeldt J, Legge R, Kim J, Sinha R, Duncan CE, Ma J, Singh I. 2014. Microbial successions are associated with changes in chemical profiles of a model refrigerated fresh pork sausage during an 80-day shelf life study. Appl Environ Microbiol 80:5178–94.

54. Jackson CR, Randolph KC, Osborn SL, Tyler HL. 2013. Culture dependent and independent analysis of bacterial communities associated with commercial salad leaf vegetables. BMC Microbiol 13:274.

55. Rice LB. 2008. Federal Funding for the Study of Antimicrobial Resistance in Nosocomial Pathogens: No ESKAPE. J Infect Dis 197:1079–1081.

56. Wang P, Wang H, Qiu Y, Ren L, Jiang B. 2018. Microbial characteristics in anaerobic digestion process of food waste for methane production–A review. Bioresour Technol 248:29–36.

57. Supaphol S, Jenkins SN, Intomo P, Waite IS, O’Donnell AG. 2011. Microbial community dynamics in mesophilic anaerobic co-digestion of mixed waste. Bioresour Technol 102:4021–4027.

58. Lin J, Zuo J, Ji R, Chen X, Liu F, Wang K, Yang Y. 2012. Methanogenic community dynamics in anaerobic co-digestion of fruit and vegetable waste and food waste. J Environ Sci 24:1288–1294.

59. Yi J, Dong B, Xue Y, Li N, Gao P, Zhao Y, Dai L, Dai X. 2014. Microbial Community Dynamics in Batch High-Solid Anaerobic Digestion of Food Waste Under Mesophilic Conditions. J Microbiol Biotechnol 24:270–279.

60. Kim YM, Jang HM, Lee K, Chantrasakdakul P, Kim D, Park KY. 2015. Changes in bacterial and archaeal communities in anaerobic digesters treating different organic wastes. Chemosphere 141:134–137.

61. Peng X, Zhang S, Li L, Zhao X, Ma Y, Shi D. 2018. Long-term high-solids anaerobic digestion of food waste: Effects of ammonia on process performance and microbial community. Bioresour Technol 262:148–158.

62. Zhang J, Zhang L, Loh K-C, Dai Y, Tong YW. 2017. Enhanced anaerobic digestion of food waste by adding activated carbon: Fate of bacterial pathogens and antibiotic resistance genes. Biochem Eng J 128:19–25.

63. Forsberg KJ, Reyes A, Wang B, Selleck EM, Sommer MOA, Dantas G. 2012. The Shared Antibiotic Resistome of Soil Bacteria and Human Pathogens. Science (80-) 337:1107–1111.

64. Nõlvak H, Truu M, Oopkaup K, Kanger K, Krustok I, Nehrenheim E, Truu J. 2018. Reduction of antibiotic resistome and integron-integrase genes in laboratory-scale photobioreactors treating municipal wastewater. Water Res 142:363–372.

65. Krawczyk PS, Lipinski L, Dziembowski A. 2018. PlasFlow: predicting plasmid sequences in metagenomic data using genome signatures. Nucleic Acids Res 46:e35.

66. Gillings MR, Gaze WH, Pruden A, Smalla K, Tiedje JM, Zhu Y-G. 2015. Using the class 1 integron-integrase gene as a proxy for anthropogenic pollution. ISME J 9:1269–79.

67. Zhang J, Chen M, Sui Q, Wang R, Tong J, Wei Y. 2016. Fate of antibiotic resistance genes and its drivers during anaerobic co-digestion of food waste and sewage sludge based on microwave pretreatment. Bioresour Technol 217:28–36.

68. O’Hara CM, Steigerwalt AG, Hill BC, Miller JM, Brenner DJ. 1998. First report of a human isolate of Erwinia persicinus. J Clin Microbiol 36:248–50.

69. Shin SY, Song JH, Ko KS. 2008. First Report of Human Infection Due to Erwinia tasmaniensis-Like Organism. Int J Infect Dis 12:e329–e330.

70. Devirgiliis C, Zinno P, Perozzi G. 2013. Update on antibiotic resistance in foodborne Lactobacillus and Lactococcus species. Front Microbiol 4:301.

71. Ju F, Zhang T. 2015. Experimental Design and Bioinformatics Analysis for the Application of Metagenomics in Environmental Sciences and Biotechnology. Environ Sci Technol 49:12628–12640.

72. Amann RI, Ludwig W, Schleifer KH. 1995. Phylogenetic identification and in situ detection of individual microbial cells without cultivation. Microbiol Mol Biol Rev 59.

73. Ferris MJ, Muyzer G, Ward DM. 1996. Denaturing gradient gel electrophoresis profiles of 16S rRNA-defined populations inhabiting a hot spring microbial mat community. Appl Environ Microbiol 62:340–6.

74. Yu Y, Lee C, Kim J, Hwang S. 2005. Group-specific primer and probe sets to detect methanogenic communities using quantitative real-time polymerase chain reaction. Biotechnol Bioeng 89:670–679.

75. Stedtfeld RD, Guo X, Stedtfeld TM, Sheng H, Williams MR, Hauschild K, Gunturu S, Tift L, Wang F, Howe A, Chai B, Yin D, Cole JR, Tiedje JM, Hashsham SA. 2018. Primer set 2.0 for highly parallel qPCR array targeting antibiotic resistance genes and mobile genetic elements. FEMS Microbiol Ecol 94:fiy130.

76. Wang F-H, Qiao M, Su J-Q, Chen Z, Zhou X, Zhu Y-G. 2014. High Throughput Profiling of Antibiotic Resistance Genes in Urban Park Soils with Reclaimed Water Irrigation. Environ Sci Technol 48:9079–9085.

77. Looft T, Johnson TA, Allen HK, Bayles DO, Alt DP, Stedtfeld RD, Sul WJ, Stedtfeld TM, Chai B, Cole JR, Hashsham SA, Tiedje JM, Stanton TB. 2012. Infeed antibiotic effects on the swine intestinal microbiome. Proc Natl Acad Sci U S A 109:1691–6.

78. Gu Z, Eils R, Schlesner M. 2016. Complex heatmaps reveal patterns and correlations in multidimensional genomic data. Bioinformatics 32:2847–2849.

79. Engelbrektson A, Kunin V, Wrighton KC, Zvenigorodsky N, Chen F, Ochman H, Hugenholtz P. 2010. Experimental factors affecting PCR-based estimates of microbial species richness and evenness. ISME J 4:642–647.

80. Caporaso JG, Kuczynski J, Stombaugh J, Bittinger K, Bushman FD, Costello EK, Fierer N, Peña AG, Goodrich JK, Gordon JI, Huttley GA, Kelley ST, Knights D, Koenig JE, Ley RE, Lozupone CA, McDonald D, Muegge BD, Pirrung M, Reeder J, Sevinsky JR, Turnbaugh PJ, Walters WA, Widmann J, Yatsunenko T, Zaneveld J, Knight R. 2010. QIIME allows analysis of high-throughput community sequencing data. Nat Methods 7:335–336.

81. Edgar RC. 2010. Search and clustering orders of magnitude faster than BLAST. Bioinformatics 26:2460–2461.

82. Bolger AM, Lohse M, Usadel B. 2014. Trimmomatic: a flexible trimmer for Illumina sequence data. Bioinformatics 30:2114–20.

83. Chen S, Huang T, Zhou Y, Han Y, Xu M, Gu J. 2017. AfterQC: automatic filtering, trimming, error removing and quality control for fastq data. BMC Bioinformatics 18:80.

84. Menzel P, Ng KL, Krogh A. 2016. Fast and sensitive taxonomic classification for metagenomics with Kaiju. Nat Commun 7:11257.

85. Bengtsson-Palme J, Hartmann M, Eriksson KM, Pal C, Thorell K, Larsson DGJ, Nilsson RH. 2015. metaxa 2: improved identification and taxonomic classification of small and large subunit rRNA in metagenomic data. Mol Ecol Resour 15:1403–1414.

86. Li D, Luo R, Liu C-M, Leung C-M, Ting H-F, Sadakane K, Yamashita H, Lam T- W. 2016. MEGAHIT v1.0: A fast and scalable metagenome assembler driven by advanced methodologies and community practices. Methods 102:3–11.

87. Nurk S, Meleshko D, Korobeynikov A, Pevzner PA. 2017. metaSPAdes: a new versatile metagenomic assembler. Genome Res 27(5):824–834.

88. Mikheenko A, Saveliev V, Gurevich A. 2016. MetaQUAST: evaluation of metagenome assemblies. Bioinformatics 32:1088–1090.

89. Wu Y-W, Simmons BA, Singer SW. 2016. MaxBin 2.0: an automated binning algorithm to recover genomes from multiple metagenomic datasets. Bioinformatics 32:605–607.

90. Kang DD, Froula J, Egan R, Wang Z. 2015. MetaBAT, an efficient tool for accurately reconstructing single genomes from complex microbial communities. PeerJ 3:e1165.

91. Olm MR, Brown CT, Brooks B, Banfield JF. 2017. dRep: a tool for fast and accurate genomic comparisons that enables improved genome recovery from metagenomes through de-replication. ISME J 11:2864–2868.

92. Parks DH, Imelfort M, Skennerton CT, Hugenholtz P, Tyson GW. 2015. CheckM: assessing the quality of microbial genomes recovered from isolates, single cells, and metagenomes. Genome Res 25:1043–55.

93. Parks DH, Rinke C, Chuvochina M, Chaumeil P-A, Woodcroft BJ, Evans PN, Hugenholtz P, Tyson GW. 2017. Recovery of nearly 8,000 metagenome-assembled genomes substantially expands the tree of life. Nat Microbiol 2:1533–1542.

94. Eren AM, Esen ÖC, Quince C, Vineis JH, Morrison HG, Sogin ML, Delmont TO. 2015. Anvi’o: an advanced analysis and visualization platform for ‘omics data. PeerJ 3:e1319.

95. Markowitz VM, Chen I-MA, Chu K, Szeto E, Palaniappan K, Pillay M, Ratner A, Huang J, Pagani I, Tringe S, Huntemann M, Billis K, Varghese N, Tennessen K, Mavromatis K, Pati A, Ivanova NN, Kyrpides NC. 2014. IMG/M 4 version of the integrated metagenome comparative analysis system. Nucleic Acids Res 42:D568–D573.

96. Shannon P, Markiel A, Ozier O, Baliga NS, Wang JT, Ramage D, Amin N, Schwikowski B, Ideker T. 2003. Cytoscape: a software environment for integrated models of biomolecular interaction networks. Genome Res 13:2498–504.

