## Supplemental Information File for "Antibiotic resistome and microbial community structure during anaerobic co-digestion of food waste, paper and cardboard"

Table of Contents

**FIG S1** Relative abundance of the most abundant bacterial and archaeal genera detected from SSU gene sequencing data and from total community metagenome sequencing data using different taxonomic annotation software.

**FIG S2** Absolute abundance of bacterial and archaeal 16S rRNA gene detected by quantitative PCR.

**FIG S3** Distribution of plasmid antibiotic resistance genes (ARGs) detected from total community metagenome sequencing data assembled with MEGAHIT.

**FIG S4** Genome statistics for 201 metagenome assembled genomes (MAGs) from 7 metagenomes (FW1, LBF1, FW2, LBF2, DG74, DG78, UASB) calculated and visualized in Anvi'o.

**Supplemental Tables**

**Supplemental Methods**

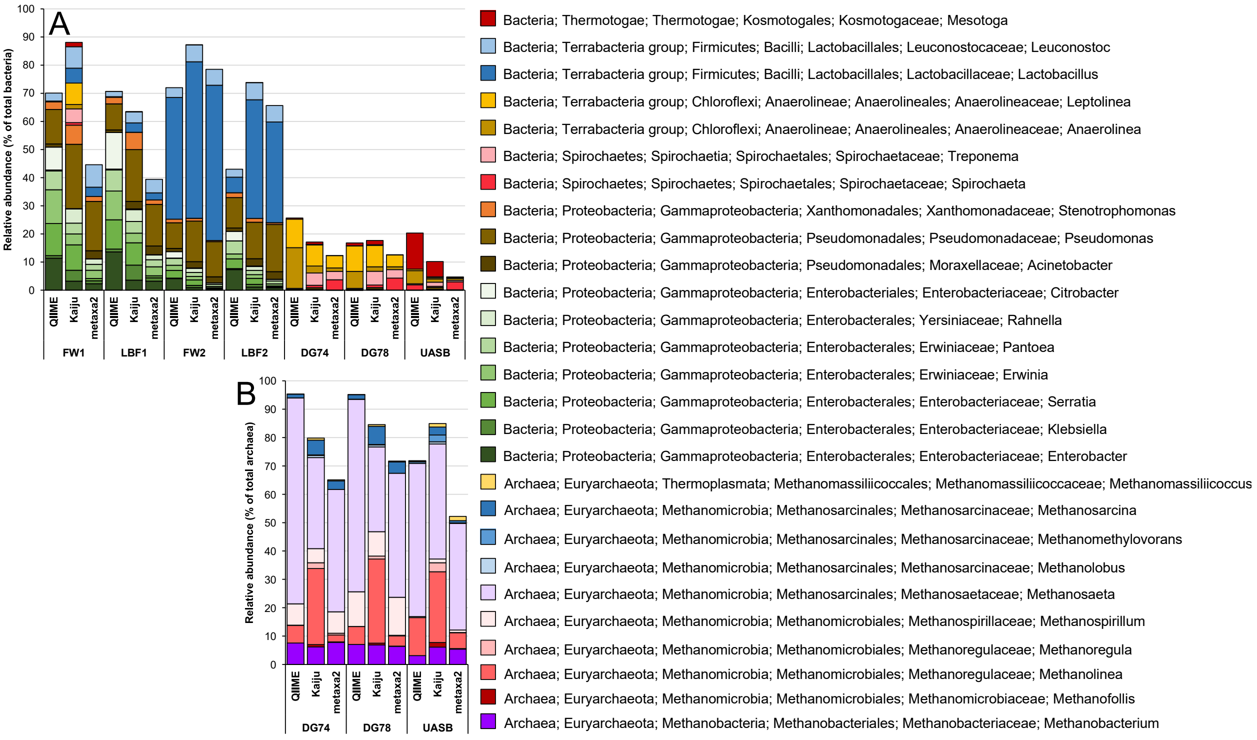


**FIG S1** Relative abundance of the most abundant bacterial (A) and archaeal (B) genera detected from SSU gene sequencing data (QIIME) and from total community metagenome sequencing data (Kaiju, metaxa2) using different taxonomic annotation software. Archaeal genera are presented only for the samples DG74, DG78, UASB due to low number of archaeal sequences in food waste (FW1, FW2) and leach bed feed (LBF1, LBF2) samples.

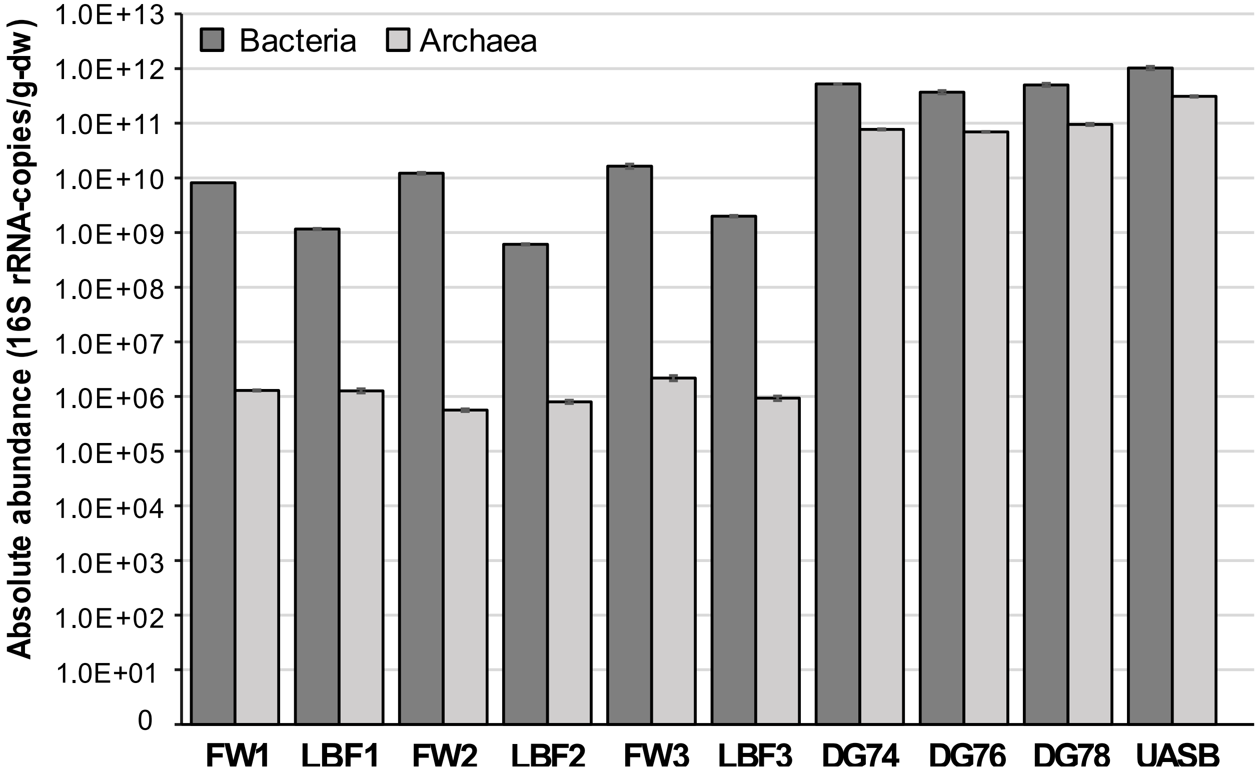


**FIG S2** Absolute abundance of bacterial and archaeal 16S rRNA gene detected by quantitative PCR. Average values with standard deviations of three technical replicates are represented on logarithmic scale.

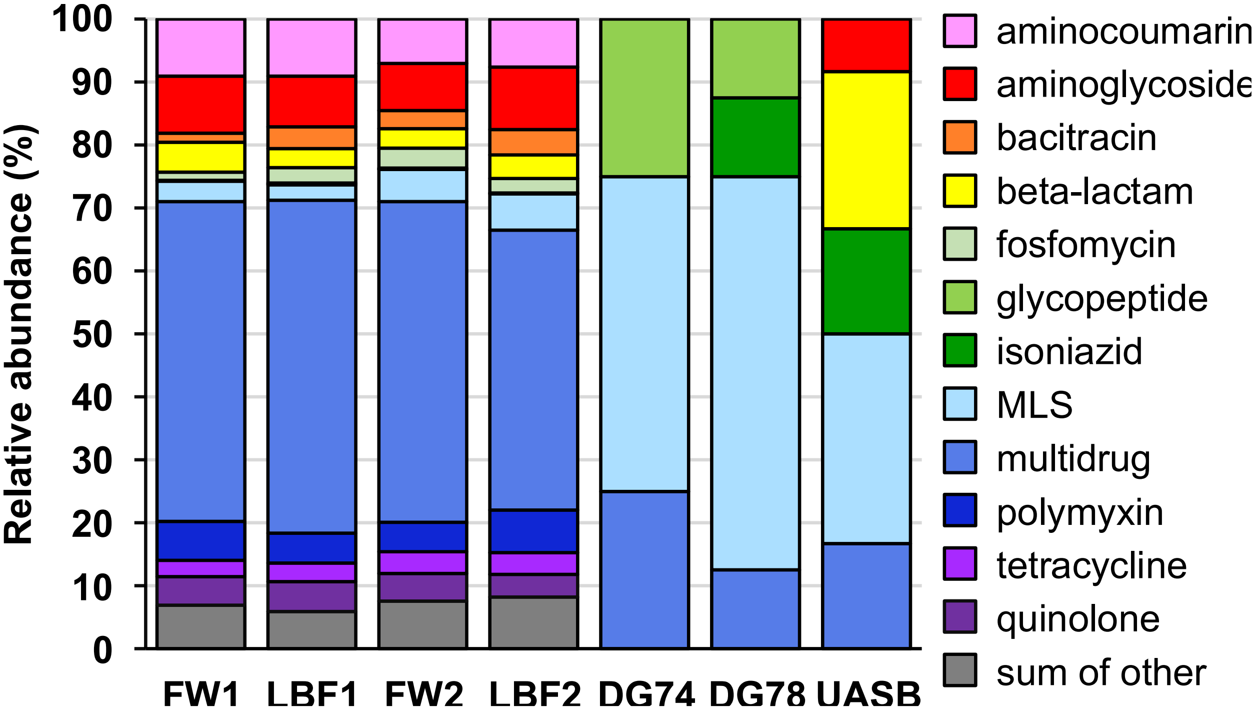


**FIG S3** Distribution of plasmid ARGs detected from total community metagenome sequencing data assembled with MEGAHIT. ARGs on contigs identified as plasmids by PlasFlow were detected by BLAST analysis of protein coding genes using local SARG reference database. Raw data for plasmid ARGs detected from total community metagenome sequencing data assembled with MEGAHIT is provided in Table S7 in supplemental material.

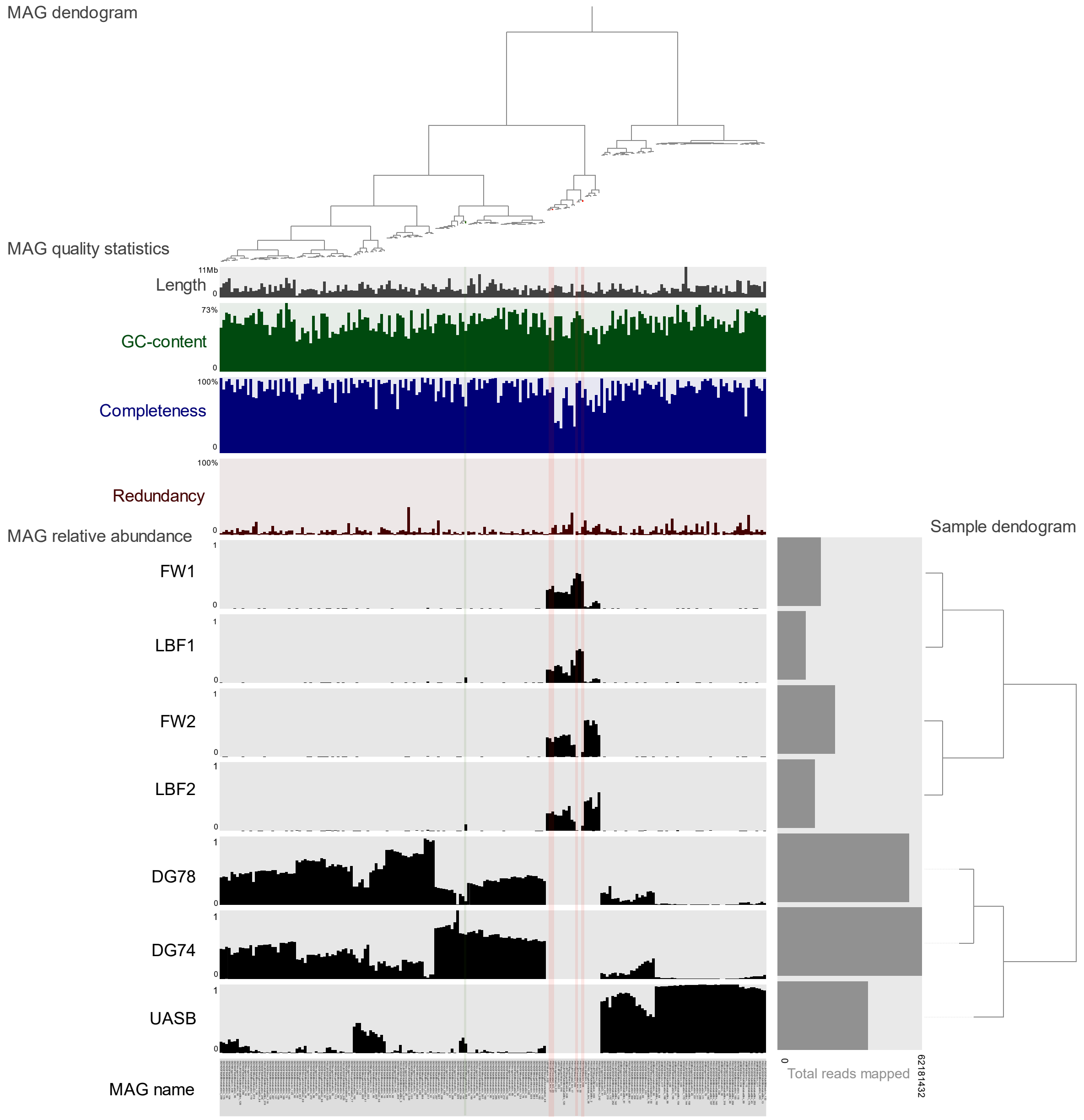


**FIG S4** Genome statistics for 201 metagenome assembled genomes (MAGs) from 7 metagenomes (FW1, LBF1, FW2, LBF2, DG74, DG78, UASB) calculated and visualized in Anvi'o. Four ARG-containing MAGs are shaded red. MAGs and samples were clustered based on the relative abundance of MAGs in the samples; Euclidian distances were calculated from relative abundance estimates of each MAG and clustering was performed using the ward linkage algorithm. Relative abundance of each MAG was calculated as number of reads recruited to the MAG divided by total reads recruited to that MAG across all samples. Length refers to total length of the MAG, and the largest MAG was a 11 Mb *Planctomycetaceae* with 1619 contigs (FW2LBF2DG78UASBmetabatBin_144). GC refers to % G + C content of the MAGs which varied between 29% and 73%. Percent completeness and redundancy were calculated based on single-copy core genes. Total number of mapped reads is shown for each metagenome sample, with the highest number of reads mapped from DG74. Bar-charts for each metagenome show the relative abundance of each MAG. Only a single MAGs was detected with relative abundance > 0.05 in both a feed and product sample (colored green): DG074DG078MegahitMaxbin.093 classified as *Porphyromonadaceae*. Additional information on all MAGs is provided in Table S9 in supplemental material.

Supplemental Tables

Supplemental tables are provided in an Excel workbook containing the tables in separate sheets.

**TABLE S1** 16S rRNA gene sequencing quality statistics.

**TABLE S2** 16S rRNA gene sequencing OTU table with OTU counts, relative abundances, reference sequences and estimated taxonomy.

**TABLE S3** Total community metagenome sequencing quality statistics.

**TABLE S4** Relative abundance of ARGs per 16S rRNA gene detected from total community metagenome sequencing reads with ARGs-OAP using default settings.

**TABLE S5** High-throughput qPCR array data for ARGs and MGEs in food waste (FW1, FW2) and digestate (DG74, DG78) samples.

**TABLE S6** Quality statistics of metagenome assembly performed with metaSPAdes and MEGAHIT assemblers.

**TABLE S7** ARGs detected from total community metagenome sequencing data assembled with MEGAHIT.

**TABLE S8** Distribution of ARGs on contigs assembled with MEGAHIT and metaSPAdes assemblers.

**TABLE S9** Metagenome assembled genomes (MAGs) from digester feed (FW1, FW2, LBF1, LBF2) and digestion products (DG74, DG78, UASB).

Supplemental Methods

Supplemental methods are publicly available online at https://github.com/kkanger/PythonScripts.

**Method S1** is a Python script to find assembled contigs with multiple antibiotic resistance genes (ARGs).

**Method S2** is a Python script to find assembled contigs that carry antibiotic resistance genes (ARGs) and are binned into metagenome assembled genomes (MAGs).
