## Supplementary material for "Antibiotic resistome and microbial community structure during anaerobic co-digestion of food waste, paper and cardboard": Method S1_python script

"""
-------------------------------------------------
Python script to find contigs with multiple ARGs
-------------------------------------------------

Author: K�rt Kanger
Date written: February 2019

This script aims at analyzing a structured tabular blast output file.
The blast output file was generated by blasting ORFs from assembled contigs against an ARG-
database.
On each line of the output file there is a blast hit, including original contig ID (first column), ORF 
ID, numerical blast results and ARG ID.
The output of this script is a text file, which shows how many contigs were found with 1 ARG, 2 
ARGs, 3 ARGs, etc, and specifies the contig IDs.

"""

# Read in the input file with tabular blast output
blastfile=open("sample_input.csv")
blast=[]
for row in blastfile:
    row = row.strip()
    row=row.split(';')
    blast.append(row)
blastfile.close()
print("Genes in blast output table: ", len(blast)-1)

# Make a list of unique contig IDs
contigs=[]
for row in blast:
    if row[0]=='Original contig ID':
        continue
    if row[0] not in contigs:
        contigs.append(row[0])
print("Number of unique contig IDs: ", len(contigs))

# Count ARG hits on each contig
results=[]
for contig in contigs:
    counter=0
    for i in range(len(blast)):
        if i==len(blast)-1: # last row
            if blast[i][0]==contig:
                counter+=1
                pair=[counter, contig]
                results.append(pair)
            else:
                pair=[counter, contig]
                results.append(pair)
        elif blast[i][0]==contig:
            counter+=1

# Make a list of number of ARGs per contig
ARG_counts=[]
for row in results:
    if row[0] not in ARG_counts:
        ARG_counts.append(row[0])
length=max(ARG_counts)

# Make a custom-sized matrix for summarizing results
summary=[]
for i in range(length+1):
    if i==0:
        summary.append(['#of ARGs per contig', '# of contigs', ['contig IDs']])
    else:
        summary.append([i, 0, []])
        
for i in range(len(summary)):
    for pair in results:
        if pair[0]==summary[i][0]:
            summary[i][1]+=1
            summary[i][2].append(pair[1])
    if i!=0:
        print("Number of contigs with ", i, " genes: ", len(summary[i][2]))

# Write summarized results to a textfile
resultsfile=open("sample_output.txt", "w")
for row in summary: 
    for i in range(len(row)): 
        if i==len(row)-1: 
            if row[i]==[]:
                resultsfile.write("\n")
            else:
                for j in range(len(row[i])): 
                    if j==len(row[i])-1: 
                        resultsfile.write(str(row[i][j])+ "\n")
                    else: 
                        resultsfile.write(str(row[i][j])+ "\t")
        else: 
            resultsfile.write(str(row[i]) + "\t")
resultsfile.close()          
