## Supplementary material for "Antibiotic resistome and microbial community structure during anaerobic co-digestion of food waste, paper and cardboard": Method S2_python script

"""
-------------------------------------------------
Python script to find ARG-contigs in MAGs
-------------------------------------------------

Author: K�rt Kanger
Date written: February 2019

This script compares a list of contig IDs that are included in metagenome assembled genomes 
(MAGs) to a list of contig IDs that carry ARGs
and outputs a list of contig IDs that carry ARGs and are included in MAGs.

Two input files are required:
1) textfile with a list of contig IDs included in the MAGs (each ID on separate line). NB: this 
script works with IDs that are formatted as "SampleName_MAGName_ContigID". In case of 
differently formatted contig IDs the script should be changed accordingly.
2) textfile with a list of contig IDs that carry ARGs (each ID on separate line).

The script outputs a textfile with a list of contig IDs that are included in MAGs and carry ARGs.

"""

# Read in the first input file with contig IDs included in the MAGs
f1=open("input_headers.txt")
headers=[]
for row in f1:
    row = row.strip()
    row=row.split('_')
    if len(row)==3:
        row=row[0]+row[1], row[2]
    else:
        print("unknown header: ", row)
    headers.append(row)
f1.close()
print("No contigs in MAGs: ", len(headers))

# Read in the second input file with contig IDs that carry ARGs
f2=open("ARGcontigs_ids.txt")
queries=[]
for row in f2:
    queries.append(row.strip())
f2.close()
print("No on ARG-contigs: ", len(queries))

# Compare ARG-contig IDs to MAG-contig IDs
results=[]
for contig in queries:
    for header in headers:
        if contig==header[1]:
            results.append(header)
print("ARG-contigs in MAGs: ", len(results))

# Write results to a textfile
f3=open("output_ARGcontigs_in_MAGs.txt", "w")
for row in results:
    for i in range(len(row)):
        if i==len(row)-1:
            f3.write(row[i]+'\n')
        else:
            f3.write(row[i]+'\t')
f3.close()    
